# Balance-directed protein engineering of *Is*PETase enhances both PET hydrolysis activity and thermostability

**DOI:** 10.1101/2022.11.14.516528

**Authors:** Seul Hoo Lee, Hogyun Seo, Hwaseok Hong, Jiyoung Park, Dongwoo Ki, Mijeong Kim, Hyung-Joon Kim, Kyung-Jin Kim

## Abstract

A mesophilic PETase from *Ideonella sakaiensis* (*Is*PETase) has been shown to exhibit high PET hydrolysis activity, but its low thermostability limits its industrial applications. We herein developed an engineering strategy for *Is*PETase to enhance PET hydrolysis activity, thermostability, and protein folding of the enzyme. Balance-directed Z1-PETase variant outperforms the stability-directed Z2-PETase variant under both mesophilic and thermophilic conditions, although Z2-PETase exhibits higher thermostability than Z1-PETase. The Z1-PETase is also superior to Fast-PETase, Dura-PETase, and LC-C^ICCG^ in terms of depolymerization rate regardless of temperature conditions we tested. Thus, maintaining a balance between PET hydrolysis activity and thermostability is essential for the development of high-performance PET hydrolases. In a pH-stat bioreactor, Z1-PETase depolymerized >90% of both transparent and colored post-consumer PET powders within 24 and 8 hours at 40°C and 55°C, respectively, demonstrating that the balance-directed *Is*PETase variant produced herein may be applicable in the bio-recycling of PET.

## Introduction

Polyethylene terephthalate (PET) is used in the production of bottles, sheets, films, and fibers that are commonly found in food packaging and clothes. It is estimated that 60 metric tons of PET are produced each year,^1^ although there are no clear estimates for forms other than solid-state PET resins. PET product wastes are largely released into the environment and known to be highly resistant to natural decomposition,^3^ making the PET industry a major contributor of plastic pollution. The major ingredients of PET, including purified terephthalic acid (pTA) and monoethylene glycol (MEG), are derived from crude oil refinery;^2^ the industry that have recourse to a petroleum-based economy is no longer sustainable despite the high demand. Petrochemical and food corporations, reclaimers, and governments are steadily seeking to improve available technologies for eco-friendly removal or closed-loop recycling of PET; however, the current methods of bottle-to-bottle mechanical recycling, pyrolysis, and chemo-depolymerization are limited^4–6^ in their usefulness.

In recent years, enzymatic PET depolymerization, which is widely applicable in the concept of a circular economy, has emerged as an alternative of the existing recycling technologies.^7^ Although no significant degradation of PET but surface modification by enzymes was reported from the mid-late 20th century to the early 2000s,^8^ it has now been shown that the depolymerization rate of PET near its glass-transition temperature (Tg) by cutinase-like enzymes is unexpectedly fast.^9–12^ In fact, Tournier *et al*. achieved 90% depolymerization of post-consumer colored-flake PET waste at 72°C in less than 10 h using an engineered variant of the thermophilic leaf-branch compost cutinase (LC-C).^12^

Yoshida *et al*. reported that the bacterium *Ideonella sakaiensis* utilizes PET as its sole source of carbon and energy via hydrolysis of PET by the *Is*PETase enzyme.^13^ This enzyme exhibits highest PET hydrolysis activity at an ambient temperature of approximately 30°C, at which the structure of PET is more glassy and difficult for enzymes to access.^13^ The mesophilic PET depolymerase brings the depolymerization system into the physiological temperature range, which hints at a possible approach for the free biotransformation of PET into any product that an organism is able to generate.^14–16^ However, low temperatures decrease the specific volume of polymeric materials, thereby increasing the thermodynamic and kinetic barriers of geometrical change, resulting in low deformability of substrates.^17^ This implies that enzyme access and complex formation with interfacial chains are impeded at low temperatures (Fig. 1a). It means that the mesophilic PET depolymerase has accessibility to overcome this limitation. In the last 6 years, several monumental thermostable and durable variants of *Is*PETase have been engineered^18–21^; however, these have not been reported to reach the stability and efficacy of thermophilic cutinases at high temperatures.

**Fig. 1.**
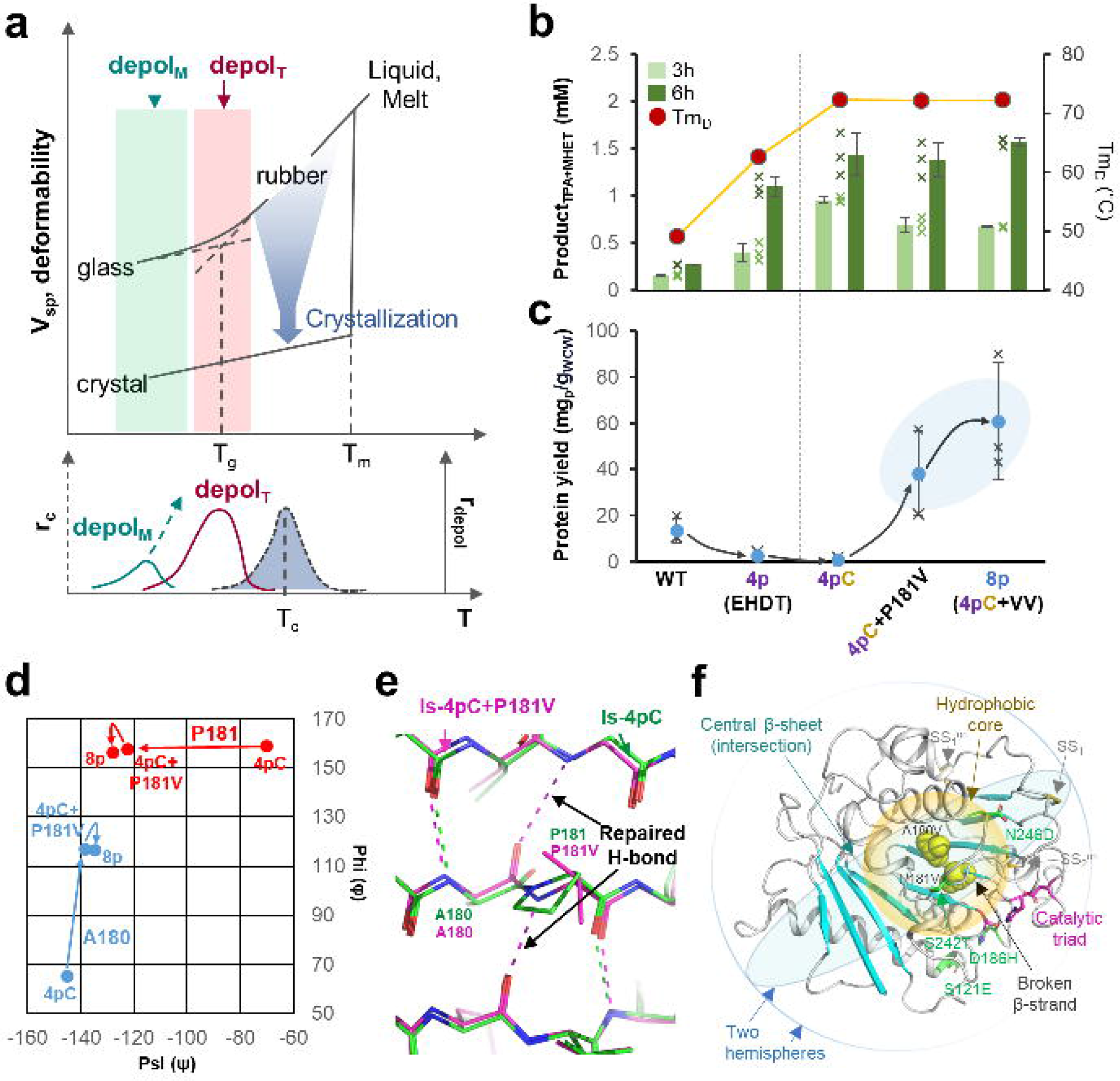
Increased protein yield through β-strand modification. (a) Schematic diagram of the relationship between temperature and PET polymer deformability. Abbreviations: V_sp_, specific volume; r_c_, crystallization rate of polymer; r_depol_, rate of depolymerization; depol_M_, mesophilic depolymerase; depol_T_, thermophilic depolymerase. (b) (c) PET hydrolysis activities and the Tm_D_ values (b) and solubilized protein yields (c) of the wild-type, Is-4p, Is-4pC + P181V, and Is-8p variants. The activity was measured at 40°C. All the experiments were performed in triplicate and detected via HPLC, and the error bars of measuring samples represent the sampling standard deviation (n=3). (d) Ramachandran plots of the A180 (blue) and P181 (red) residues from Is-4pC, Is-4pC+P181V, and Is-8p. (e) The core structures of Is-4pC (green) and Is-4pC+P181V (magenta). The polar β-strand network is shown with dotted lines. (f) The hydrophobic core and the variations. The crystal structure of Is-8p is shown with a gray cartoon model. The catalytic triad of Is-8p is distinguished with magenta color and their side chains are shown with sticks. The mutation points of 4p, SS_1_, A180V, and P181V are indicated and labeled. The two imaginary hemispheres and the hydrophobic core of the protein are shown with light-blue and yellow circles, respectively.

In this work, we present our engineered PET depolymerase (Z1-PETase) that exhibits high thermostability, PET hydrolysis activity, and soluble protein yield. Z1-PETase outperforms the stability-directed Z2-PETase variant, Dura-PETase, LC-C^ICCG^, and Fast-PETase variants over a broad range of temperatures (30°C–60°C) in a dialysis system. Z1-PETase showed a decomposition rate of >90% for post-consumer PET powder within 24 h at 40°C and 8 h at 55°C in a pH-stat bioreactor. These results emphasize the importance of maintaining a balance between enzyme activity and thermostability in the engineering of PET hydrolases.

## Results

### Increase in protein yield through β-strand modification of the hydrophobic core

Because amorphous polymeric chains can be crystallized above Tg and the deformability of the crystalline region is much lower than that of the amorphous region, the optimal temperature for the operation of an artificial enzyme depolymerization system is approximately 60°C–70°C (Fig. 1a).^12,23^ We hypothesized that a mesophilic depolymerase emerged to have improved thermal stability would outperform other thermophilic enzymes at all temperatures, as high activity at low temperatures is a result of the sum effect of turnover rate, adsorption rate, and attack capacity on solid PET (Fig. 1a). Thus, we evaluated the potential activity of a mesophilic depolymerase at desired temperatures with an aim of improving the thermal stability of *Is*PETase without compromising its activity. To this end, we introduced N233C and S282C mutations to create an additional disulfide bond (SS_1_) in the quadruple-point *Is*PETase^S121E/D186H/S242T/N246D^ variant (Is-4p) that we previously reported,^22^ resulting in the generation of the *Is*PETase^S121E/D186H/S242T/N233C/N246D/S288C^ variant (Is-4pC) (Supplementary Tables 1 and 2).^22,24^ Surprisingly, the derivative melting temperature (Tm_D_) of Is-4pC was found to be 72°C, which was 10°C and 23°C higher than that of Is-4p and wild-type *Is*PETase, respectively (Fig. 1b and Supplementary Fig. 1). Moreover, Is-4pC exhibited 1.5 and 5-fold higher PET depolymerization activity than those of Is-4p and wild-type *Is*PETase, respectively, at 40°C (Fig. 1b). However, the expression of Is-4pC dramatically reduced the amount of soluble protein compared with wild-type, and most of the expressed protein formed insoluble aggregates (Fig. 1c, Supplementary Figs. 2 and 3). This limited further evaluation of PET depolymerization capacity and analysis of potential applications of the variant. The production of large amounts of soluble PET hydrolase is an important parameter in industrial-level PET biodegradation in addition to its thermal stability and activity. Thus, we attempted to increase the solubility of *Is*PETase by reconstructing the β-sheet that is partially disrupted by P181^18^, replaced P181 with various other residues, including A, G, S, C, T, V, I, L, Y, and F in the Is-4p template. Interestingly, all variants exhibited up to 25-fold increase in soluble protein yield compared with Is-4p (Fig. 1c, Supplementary Figs. 2 and 3). However, most variants showed a dramatically decrease in Tm_D_ values and PET hydrolysis activities (Supplementary Figs. 1 and 3). One exception was the P181V variant, which exhibited comparable Tm_D_ value and PET hydrolysis activity to Is-4p (Fig. 1c, Supplementary Fig. 3). Introduction of the P181V mutation into Is-4pC resulted in a 40-fold increase in soluble protein yield compared with Is-4pC, with comparable Tm_D_ value and PET hydrolysis activity (Fig. 1b, c, Supplementary Figs. 1–3). We further observed no significant difference between Is-4pC+P181V and Is-4pC in Tm_D_ values and the total amount of expressed protein, which indicate that the enhanced yield of the soluble Is-4pC+P181V protein is not due to increased intracellular protein stability or enhanced transcriptional and translational processes, but rather due to post-translational folding. Specifically, mutation of the trans-proline 181 unchains the dihedral angles of this residue, especially at the ψ angle, triggering a large change in the φ angle of the pre-proline A180 (Fig. 1d). We speculated that this structural change occurs in most P181-point variants and strengthens the β-strand network, thereby increasing the efficiency of folding of the Is-4pC+P181V variant (Fig. 1d, e and Supplementary Fig. 4).

The region containing A180 and P181 is located at the intersection of two imaginary hemispheres of the protein hydrophobic core. The above results indicate the importance of this region in folding of the protein (Fig. 1f). We replaced A180 of Is-4pC+P181V with other small residues, such as G, S, T, and V, to investigate how further modification of the region affects protein solubility. Interestingly, all four variants showed an increase in soluble protein yield (Fig. 1c and Supplementary Figs. 2 and 3). Notably, Is-4pC+A180V+P181V (an Is-8p variant) showed an approximately 65% increase in soluble protein yield compared with Is-4pC+P181V, which corresponds to a 65-fold increase compared with Is-4pC (Fig. 1c, and Supplementary Figs. 2 and 3). Moreover, the thermal stability and PET hydrolysis activity of Is-8p were comparable to those of Is-4pC (Fig. 1b and Supplementary Figs. 1 and 3). This implies that the folding process of *Is*PETase is influenced by multiple structural factors as well as β-sheet repair at the protein core.

### Engineering with non-negative and complementary accumulation

The improved soluble protein yield of Is-8p led us to apply various enzyme engineering approaches. We modified the local surface charge of Is-8p by mutating R132 and R224 to E and substituting N37 with D in an attempt to optimize the adsorption process of the protein to the PET substrate. Although the Tm_D_ values of the R132E and R224E mutants were slightly lower than that of Is-8p, all three variants showed up to 50% enhancement in PET hydrolysis activity at 50°C compared with Is-8p (Fig. 2 and Supplementary Fig. 1). To further enhance the thermal stability of Is-8p, we introduced six new disulfide bonds by creating the following double mutants: A171C+S193C (SS_2_), A202C+V211C (SS_3_), N73C+A102C (SS_4_), A80C+V149C (SS_5_), T195C+R224C (SS_6_), and N275C+F284C (SS_7_). Among these, SS_4_, SS_5_, and SS_6_ had decreased Tm_D_ values compared with Is-8p, implying that disulfide bonds were not formed in these mutants (Fig. 2a and Supplementary Fig. 1). However, SS_2_, SS_3_, and SS_7_ increased Tm_D_ values by approximately 4°C, suggesting that these disulfide bonds were indeed present in Is-8p (Fig. 2a, Supplementary Fig. 1). The PET hydrolysis activity of the SS_2_ mutation increased by 20%, whereas that of SS_3_ and SS_7_ mutations slightly decreased compared with Is-8p at 50°C (Fig. 2a). The wobbling W185 residue is known to be crucial for both enzyme activity and protein stability.^26^ Because S214 is located in the vicinity of this residue, we replaced S214 with 19 other residues to identify the optimal residue in terms of enzyme activity and protein stability. Although PET hydrolysis activity decreased compared with that of Is-8p in all cases, thermal stability was improved in several of the new variants (Supplementary Figs. 1 and 5). In particular, the Tm_D_ value of S214Y was 6.5°C higher than that of Is-8p (Fig. 2a).

**Fig. 2.**
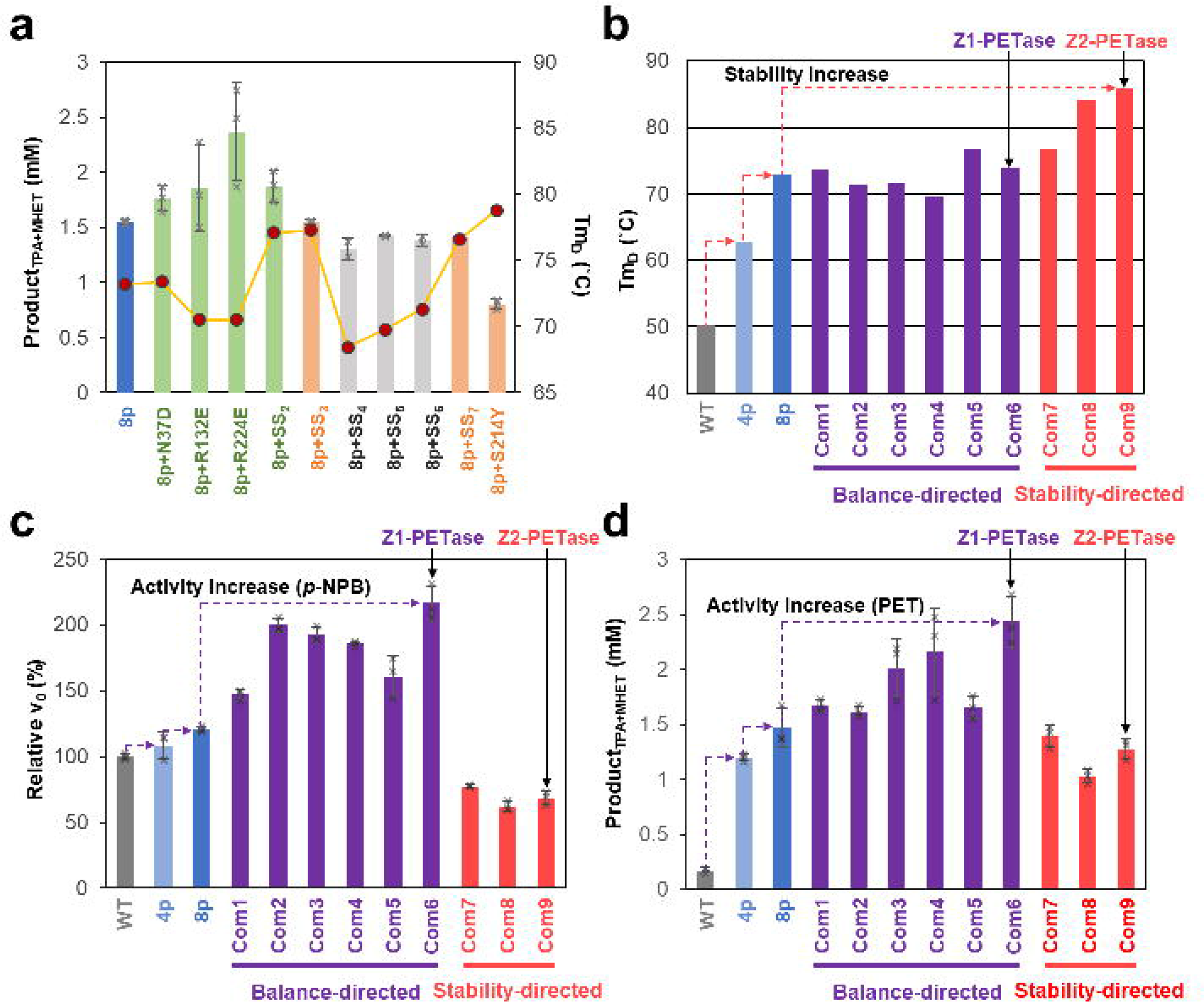
Balance- and stability-directed protein engineering. (a) PET hydrolysis activities of the mutations added to Is-8p for 6 h at 50°C. (b-d) The Tm_D_ values (b), relative activity against *p*-NPB (c), and PET hydrolysis activity (d) of the nine combinatorial variants of 8p. All the experiments were performed in triplicate and the error bars of measuring samples represent the sampling standard deviation (n=3).

Modification of Is-8p enabled us to identify seven mutation points that improved thermal stability, enhanced PET hydrolysis activity, or had other complementary effects. Next, we combined multiple mutations to design a total of nine combinatorial variants of Is-8p, including six balance-directed variants (Is-8p^Com1^–Is-8p^Com6^) and three stability-directed variants (Is-8p^Com7^–Is-8p^Com9^) (Table 1). All six balance-directed variants exhibited enhanced activities against both *p*-nitrophenyl butyrate (*p*-NPB) and PET, along with either increased (Is-8p^Com1^, Is-8p^Com5^, and Is-8p^Com6^) or decreased (Is-8p^Com2^, Is-8p^Com3^, and Is-8p^Com4^) Tm_D_ values compared with Is-8p (Fig. 2b–d and Supplementary Fig. 1). All three stability-directed variants increased Tm_D_ values compared with Is-8p but decreased enzyme activity (Fig. 2b–d and Supplementary Fig. 1). These results indicate that the combinatorial variants were generated in the intended directions, and we selected the Is-8p^Com6^ (Z1-PETase) and Is-8p^Com9^ (Z2-PETase) variants as the final balance- and stability-directed variants, respectively (Fig. 2b – d). Z2-PETase with a Tm_D_ value of 86°C has much higher thermostability than Z1-PETase with a Tm_D_ value of 74°C; conversely, Z1-PETase showed a 50% increase in PET hydrolysis activity while Z2-PETase showed somewhat decreased activity, compared with Is-8p (Fig. 2b, d).

**Table 1.**
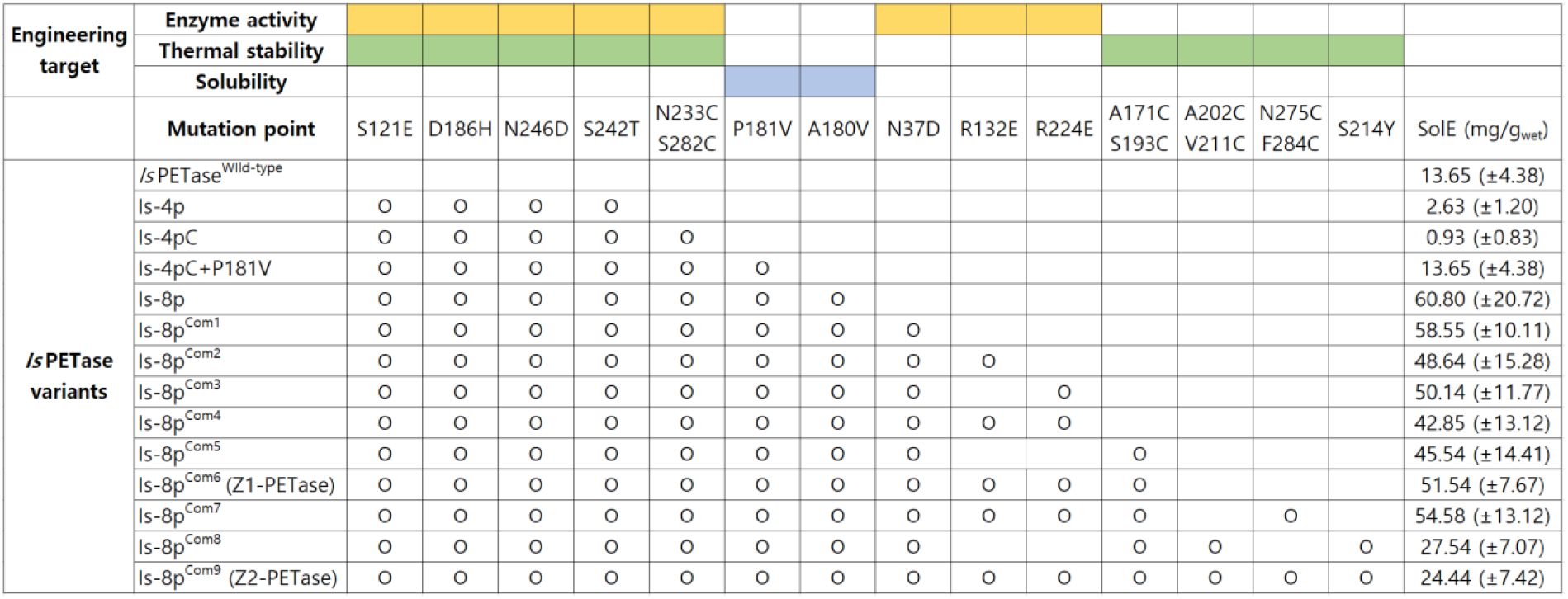
Mutation points used applied to the *Is*PETase combinatorial variants in this study. The engineering target was classified into three effects; enzyme activity, terminal stability, and solubility were marked on top in yellow, green, and blue, respectively. The soluble expression (SolE, mg/g_wet_) of combinatorial *Is*PETase variants are cultured and purified in triplicate and the error ranges represent the sampling standard deviation (n=3).

### Structural interpretation of the engineered mutants

The crystal structure of Z2-PETase provides a structural explanation of the effects of the introduced mutations in this study (Supplementary Table. 3). We identified two disulfide bonds of *Is*PETase wild-type, i.e., SS_1_^Ori^ and SS_2_^Ori^, in Z2-PETase (Fig. 3). The four mutated residues in Is-4p, namely, S121E, D186H, S242T, and N246D, were found to be located at identical positions as those in Z2-PETase (Fig. 3 and Table. 1). All Is-8p^Com^ variants having the A180V and P181V mutations in Z2-PETase showed a significant increase in soluble protein production level compared with Is-8p, although the increase in production level was somewhat reduced by further addition of mutation points (Table. 1 and Supplementary Fig. 6). Three mutations, i.e., N37D, R132E, and R224E, resulted in a large change in the protein surface charge to negative, and in fact, the pI values of wild-type and Z2-PETase were 9.44 and 7.54, respectively. The charge change occurs far from the active site, likely facilitating the adsorption of the enzyme to the PET substrate and thereby increasing PET hydrolysis activity (Fig. 3 and Supplementary Fig. 7). The four new disulfide bonds (SS_1_, SS_2_, SS_3_, and SS_7_) that were formed after the introduction of mutations stabilized the periphery of each disulfide-formed region in Z2-PETase (Fig. 3 and Supplementary Figs. 8 and 9). Although the length of SS_3_ was 2.3 Å, which was slightly longer than the optimal length of a disulfide bond, the other three disulfide bonds were within the optimal disulfide bond length range (Supplementary Fig. 8). The S214Y mutation stabilized the region surrounding the wobbling W185; however, the S214Y mutation appears to interfere with the dynamics of W185, resulting in a dramatic decrease in PET hydrolysis activity (Fig. 3 and Supplementary Fig. 9).

**Fig. 3.**
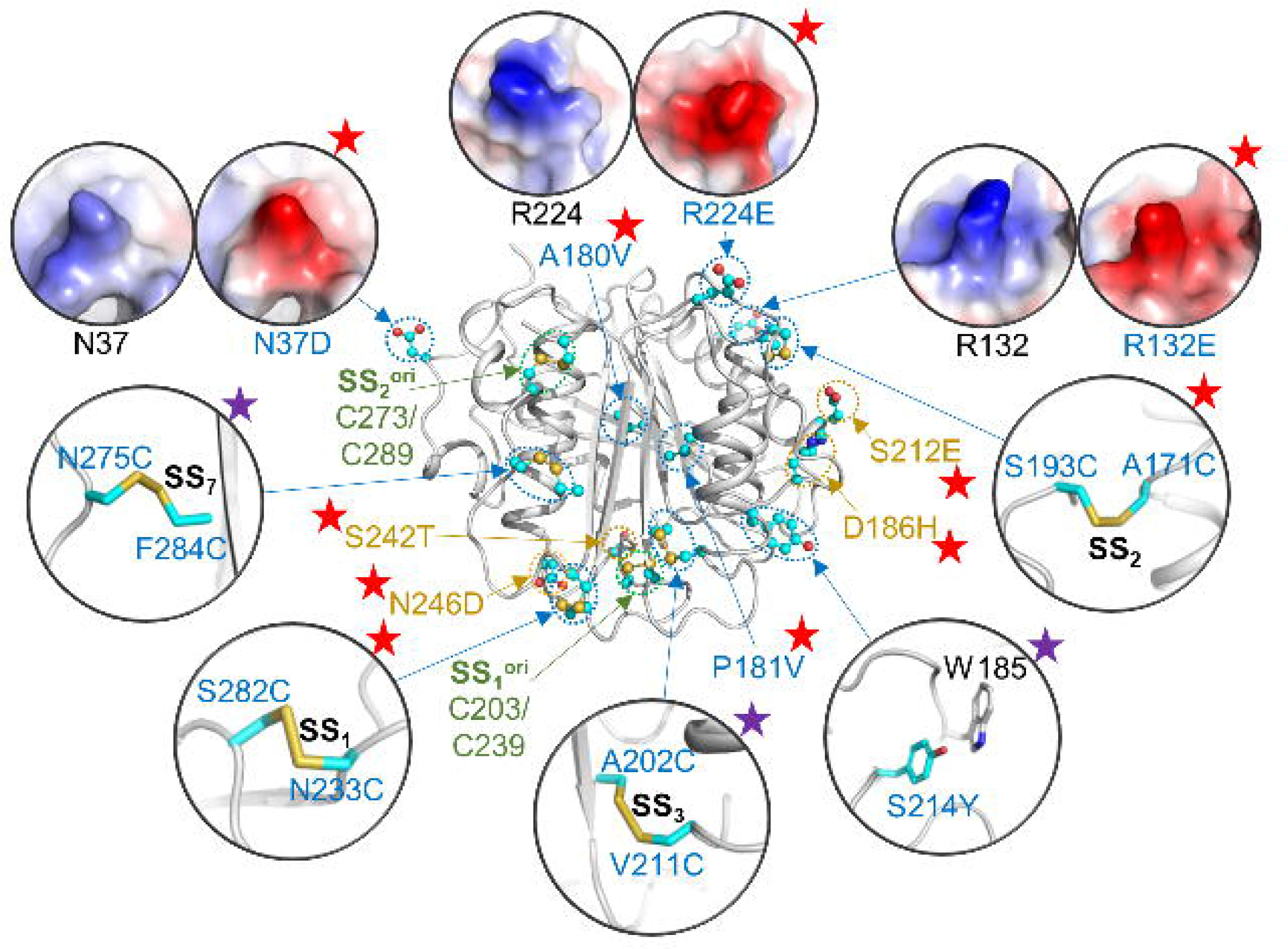
Structural interpretation of the introduced mutations. The Z2-PETase structure is shown as a gray cartoon model. The mutated residues are shown as a cyan-colored ball-and-stick model. The mutation points applied to Z1-PETase are marked with red stars, and the additional mutation points applied to Z2-PETase are with purple stars.

### Evaluation of variants in a product-deficient system

We compared Z1-PETase and Z2-PETase to identify the most superb *Is*PETase variant. The stability-directed variant Z2-PETase was much more thermostable than the balance-directed variant Z1-PETase; ΔTm_D_ between the two variants was found to be 12°C (Fig. 2b). The results of heat inactivation experiments verified the high thermostability of Z2-PETase as activity was maintained for approximately 10 h at 70°C, whereas Z1-PETase activity was lost after 1 h (Fig. 4a). This further confirms the increased thermostability of Z2-PETase compared with Z1-PETase. However, Z1-PETase maintained its activity for several days at 60°C, indicating somewhat higher thermostability than Is-8p (Fig. 4a).

**Fig. 4.**
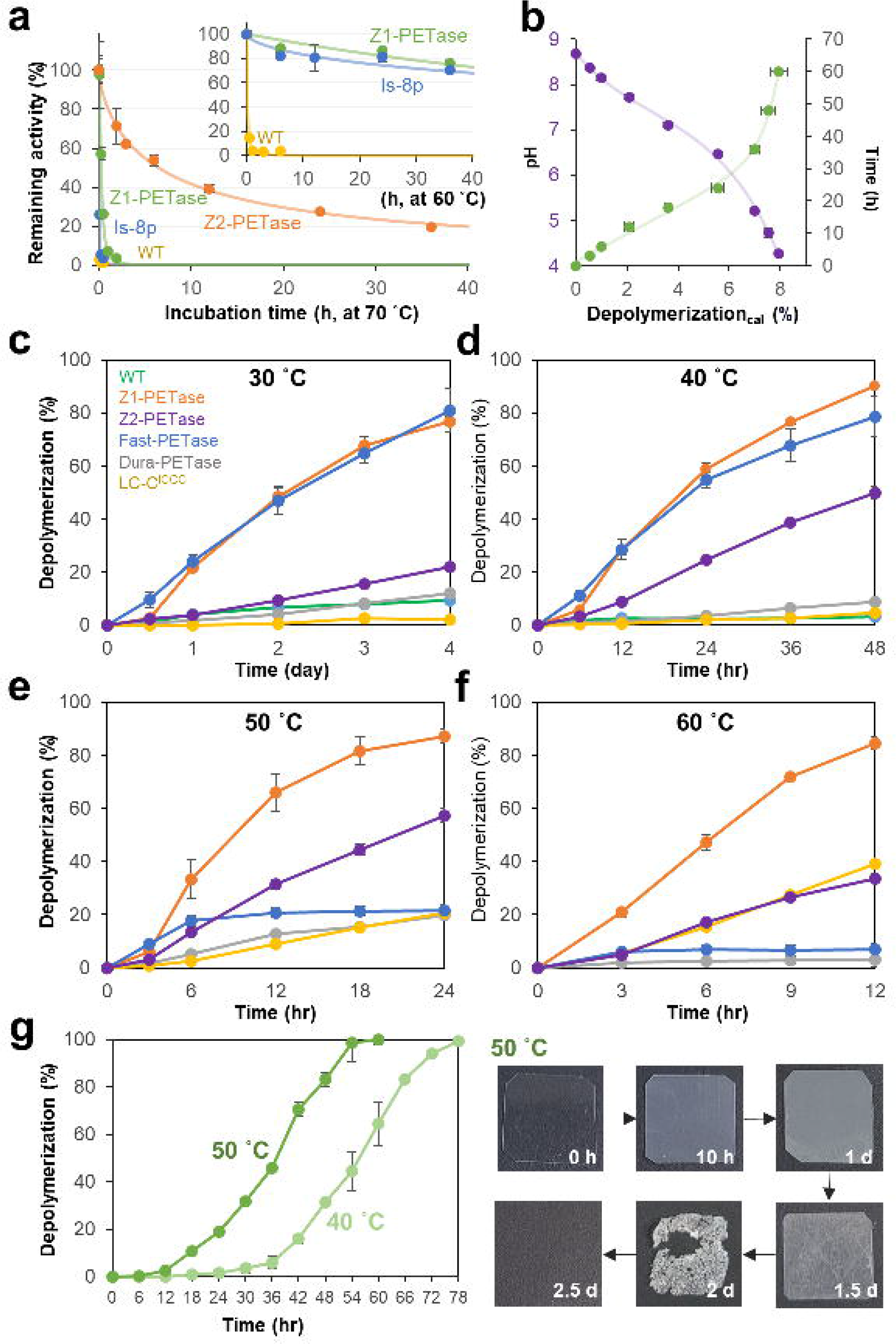
Comparison of PET depolymerization activity of Z1-PETase with those of other engineered PET hydrolases. (a) Heat inactivation assay of Z1-PETase and Z2-PETase. (b) Time-course change of pH and depolymerization rate. (c-f) Comparison of PET depolymerization activity of Z1-PETase with those of other engineered PET hydrolases. (g) Depolymerization of PET film by Z1-PETase at 40°C and 50°C. All the experiments were performed in triplicate, and the error bars of measuring samples represent the sampling standard deviation (n=3).

The closed PET hydrolysis system used above has considerable limitations for accurate quantification of enzyme kinetic performance and evaluation of complete PET depolymerization because continuous ester hydrolysis interferes with enzyme activity by lowering the pH of the reaction mixture and the inhibition of the enzyme activity by the hydrolysis products (Fig. 4b). Thus, to compare PET hydrolysis by Z1-PETase and Z2-PETase over a broad range of temperatures, we designed a product-deficient *in*-*situ* dialysis system to maintain the pH at 8.0 and keep the level of hydrolysis products low. This system was applied to PET hydrolysis (Supplementary Fig. 10), which showed that the PET depolymerizing activity of Z1-PETase was higher than that of Z2-PETase across the entire temperature range from 30°C to 60°C, although the optimal temperatures of these two variants were different (Fig. 4c–f). These results suggest that maintaining or enhancing the high activity of mesophilic PET hydrolase is critical for the development of a high-performance PET hydrolase with enhanced thermostability, and consequentially, we selected Z1-PETase as the *Is*PETase variant with the highest PET depolymerizing activity among the tested combinatorial variants.

We then compared the PET depolymerizing activity of Z1-PETase with those of other engineered PET hydrolases such as LC-C^ICCG^, Dura-PETase, and Fast-PETase, and surprisingly, Z1-PETase kinetically outperforms LC-C^ICCG^ and Dura-PETase at all temperatures tested and exhibits high depolymerization rates (76.9%–90.1%) (Fig. 4c–f). Fast-PETase exhibited a slightly faster lag phase than Z1-PETase at 30°C and 40°C; however, activity decreased dramatically at 50°C and 60°C, which was consistent with the much lower thermostability of Fast-PETase (64°C)^20^ than Z1-PETase (Fig. 4c-f). Interestingly, the PET depolymerizing activity of Dura-PETase was much lower than that of Z1-PETase at all temperature ranges (Fig. 4c-f), although the thermostability of these *Is*PETase variants was similar. These results further emphasize the importance of maintaining or enhancing mesophilic PET hydrolysis activity during the development of high-performance PET hydrolases.

Z1-PETase was able to completely depolymerize the 15 × 15 × 0.25-mm PET film (6.1% crystallinity) at 40°C and 50°C within approximately 3.2 and 2.2 days, respectively (Fig. 4g). Film decomposition showed a clear lag phase, which reflects the time period required for the formation of a sufficient contact area between the surface of PET and the enzyme via initial hydrophilization of the film (Fig. 4g). The lag phase of film decomposition was much longer than that of powder decomposition at the same temperature (Fig. 4 d, e, g), suggesting that the availability of the PET surface for enzyme adsorption determines the initial depolymerization rate.

### Depolymerization of post-consumer PET powder using a pH-stat bioreactor

We evaluated the industrial application of Z1-PETase by studying PET plastic depolymerization using a pH-stat bioreactor. We used transparent post-consumer PET powder originated from transparent PET flakes collected in Daegu, Korea as the substrate (Fig. 5a and Supplementary Fig. 11). When the bioreactor was operated at 40°C under the pH-8.0 stationary condition with 2.5% (w/v) of the substrate, 50% of the PET powder was depolymerized in 8 h, and 92.5% of the PET powder was depolymerized in 24 h (Fig. 5b). At 55°C, the initial depolymerization rate was much higher than that at 40°C, demonstrating the thermal effect on PET degradation. At 55°C, 50% and 93% of the PET powder were depolymerized in 2 h and 8 h, respectively (Fig. 5b). This corresponds to a specific productivity value of 5.2 g_monomer_∙l^−1^∙h^−1^∙μmol_enzyme_^−1^, with a maximum rate of 17.2 g_monomer_∙l^−1^∙h^−1^∙μmol_enzyme_ ^−1^ observed in the early phase of the reaction. Due to the high activity and stability of Z1-PETase, the final depolymerization rates of PET powder at 40°C and 55°C were not significantly different from each other (Fig. 5b).

**Fig. 5.**
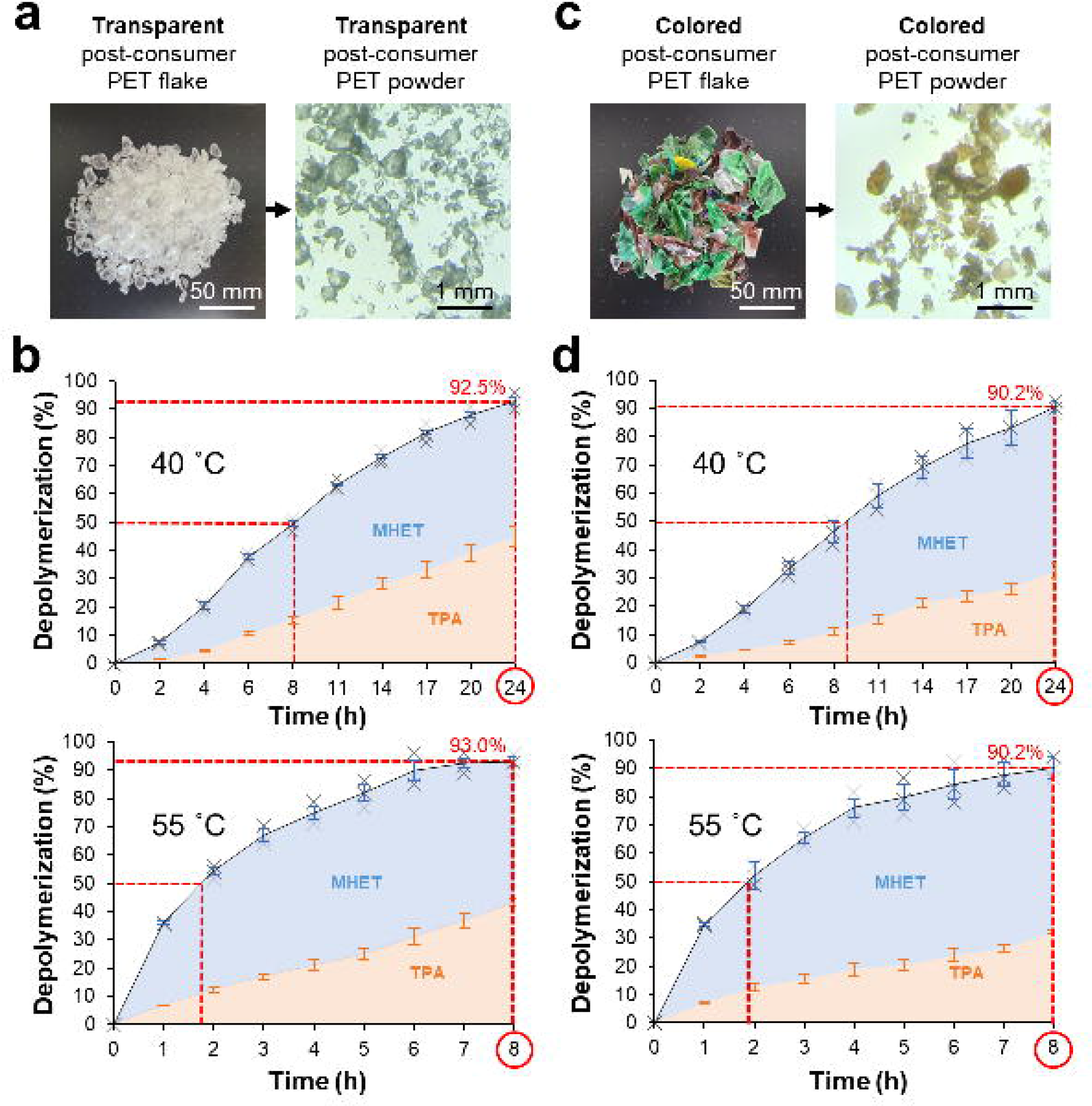
Post-consumer PET depolymerization by Z1-PETase in a pH-stat bioreactor. (a) The transparent PET flake (left) and its pulverized form (right). (b) Depolymerization of post-consumer transparent PET powder using Z1-PETase in a pH-stat bioreactor with a 3mg_enzyme_∙g ^−1^. (c) The colored PET flake (left) and its pulverized form (right). (d) Depolymerization of post-consumer colored PET powder using Z1-PETase in a pH-stat bioreactor with a 3mg_enzyme_∙g ^−1^. All the experiments were performed in triplicate, and the error bars of measuring samples represent the sampling standard deviation (n=3).

It should be noted that colored PET plastics, such as brown plastic beer bottles, are difficult to recycle because they contain impurities, such as pigments, nylon and iron. Therefore, colored PET plastics reduce the quality of transparent PET plastics when both are mixed, and colored PET plastics are not suitable as raw materials for applications in the textile industry. We prepared colored post-consumer PET powder in the same way as transparent post-consumer PET powder and performed decomposition experiments using a pH-stat bioreactor to test the efficacy of our system for difficult-to-recycle substrates (Fig. 5c and Supplementary Fig. 11). Interestingly, Z1-PETase was able to depolymerize colored PET at a rate similar to that of transparent PET at both 40°C and 55°C, indicating that Z1-PETase can depolymerize at the same rate regardless of the types of PET plastics (Fig. 5b, d). At both temperatures, the final depolymerization rate of colored PET was 90.2%, which was slightly lower than that of transparent PET, possibly due to impurities contained in colored PET (Fig. 5d).

### Discussion

Proteins are classified as psychrophilic, mesophilic, or thermophilic based on their optimal temperature, and the temperature-based classification has also been applied to depolymerases. Active site flexibility is frequently observed in psychrophilic proteins; this makes enzymes more heat-labile but decreases the temperature dependence of catalysis by reducing activation enthalpy compensated with more negative activation entropy, which does not improve the overall catalytic rate of enzyme.^25^ In contrast, high accessibility of enzyme increases the likelihood of complexation at attackable sites on PET surface, even at high temperatures at which substrate conformation is still constrained. We hypothesized that the mesophilic behavior of *Is*PETase is primarily due to its good accessibility combined with heat lability and that the reaction rate could therefore be maximized if both activity and stability are optimized; but these properties could be conflicting in the mechanism of cold-adapted enzymes. The soluble protein yield is an aspect of protein production that is critical for application at an industrial level. Thus, we simultaneously considered three crucial factors, namely, enzyme activity, thermostability, and protein folding, to develop a potent PET hydrolase variant and designed a balance-directed protein engineering strategy through non-negative and complementary accumulation. The balance-directed variant Z1-PETase and the stability-directed variant Z2-PETase highlight the importance of considering the above criteria for developing and evaluating enzymes suitable for creating an efficient PET degradation system.

## Methods

### Site-directed mutagenesis

The template used to create all variants is shown in the Supplementary Table. 2.^22^ The variant *Is*PETase^S121E/D186H/S242T/N246D^ was subcloned into a pET-15b expression vector at the NdeI and XhoI restriction sites. The forward and reverse primers used are detailed in Supplementary Table. 1. The polymerase chain reaction (PCR) step of site-directed mutagenesis was performed using EzPCR™ (EBT-7751, ELPIS-BIOTECH, Daejeon, Republic of Korea), and site-directed mutagenesis was performed using the QuikChange Site-Directed Mutagenesis Kit (Agilent). The introduction of mutations was confirmed through sequencing, which was performed by Bioneer (Daejeon, Republic of Korea).

### Protein preparation

The pET-15b expression vectors containing the desired genes were transformed into *Escherichia coli* Rosetta gami-B cells. Cells were cultured at 37°C in 1 L of lysogeny broth medium containing 200 mg L^−1^ ampicillin until the OD600 reached 0.6–7. To induce protein overexpression, 500 μM isopropyl β-D-1-thiogalactopyranoside (IPTG) was added and the cultures were further incubated for 22 h at 18°C. Cells were then harvested via centrifugation (4,000 × g for 15 min), and the obtained cell pellet was sonicated using buffer A (50 mM Na_2_HPO_4_-HCl, pH 7.0). Cell debris was removed via centrifugation (13K × g for 30 min). Purification of all proteins was performed using a Ni-NTA agarose column (Qiagen). After washing with buffer A containing 30 mM imidazole, bound proteins were eluted with 15 mL of buffer B (50 mM Na_2_HPO_4_-HCl, pH 7.0, 300 mM imidazole). All procedures were conducted at 4°C. Protein quantification was performed using a BioTek™ Epoch microplate spectrophotometer and Gen5™ microplate data analysis software.

### Differential scanning fluorometry

A thermal shift assay was performed to evaluate the thermal stability of *Is*PETase variants via StepOnePlus Real-Time PCR (Thermo Fisher Scientific). Protein Applied Biosystem 1× dye, buffer A, and 5 μg of protein were mixed in a final volume of 20 μl. Signal changes reflecting protein denaturation were monitored while the temperature increased from 25°C to 99°C at a rate of 0.1°C∙s^−1^. The melting temperature was measured by generating a derivative curve from the fluorescence melting curves of the protein complex with the Applied Biosystems dye.

### PET sample preparation

Bottle-derivative PET powder was prepared by pulverizing clear transparent bottles using a document crusher and melting the pulverized 0.2 × 1.0-mm flakes at 270°C for 7–8 min in a high-temperature oven. Molten PET was instantly chilled and hardened by immersion in a water bath set at 4°C. The obtained product was treated with cryogenic grinding (freezing milling) again and then sieved using a mesh size of 300 μm to obtain the final bottle-derived PET powder. Transparent and colored post-consumer PET powders produced from PET products that had been collected and separated into transparent and colored forms were purchased from Fluent R&C company (Deagu, Republic of Korea; 1 ton of each). Each sample was re-ground in a frozen powder dummy after generating chips by re-grinding and extrusion at high temperature and high pressure. Each sample was sieved using a mesh size of 300 μm and processed in Daehan Industrial Co., Ltd. (Deagu, Republic of Korea).

### Crystallinity measurement of PET samples

The percentage of crystallinity (Xc) for each PET sample was measured using a differential scanning calorimetry instrument (Q2000, TA instrument). Approximately 5–7 mg of each powder sample was prepared and equilibrated at 30°C. The sample was temperature scanned from 30°C to 300°C at a controlled constant rate (e.g., 10°C∙min^−1^) with heat-flow monitoring to characterize the thermal properties as a function of temperature change. Results were plotted as graphs with temperature (°C) on the x-axis and heat flow (W/g) on the y-axis, and the percentage of crystallinity was calculated based on the following equation:

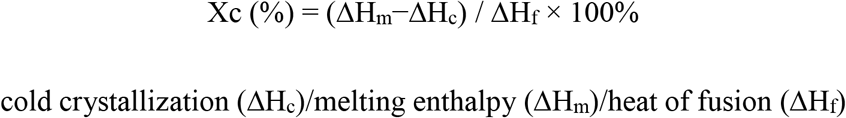

PET film from Goodfellow (thickness: 0.25 mm; condition: transparent, amorphous) is known to have a crystallinity of 6.1%.^27^

### PET hydrolysis activity measurement

The 15g ∙ l^−1^ bottle-derived PET powder was soaked in 1000 μl of 50 mM glycine/NaOH, pH 8.7, with 500 nM of *Is*PETase variants. The reaction mixture was incubated at 50°C for 3, 6, or 12 h. Then, the reaction was terminated by heating the mixture at 100°C for 15 min. The remaining substrate was removed from the reaction mixture by filtering with a 0.22-μm polyvinylidene fluoride (PVDF) syringe filter (SFPVDF013022CB, Futecs, Republic of Korea). The supernatant was analyzed using high-performance liquid chromatography (HPLC). To measure PET decomposition in a circulating system, an experimental container (350 ml) capable of accommodating a capacity of 50× the reaction volume (200 ml) was prepared. The PET hydrolase reaction was induced on a cellulose dialysis membrane (Spectra/Por 2 Trial Kit, 12–14 kD: flat width 25 mm; Repligen, USA), and bottle-derived PET powder was used as the substrate. Then, 1 μM of each variant was added to a solution of 50 (±2) mg of the substrate as bottle-derivative PET powder, 200 mM Tris-HCl (pH 9.0), and 1% (v/v) 1-butanol. Each packaged 4-ml dialysis reaction bag was immersed in 200 ml of 200 mM Tris-HCl (pH 9.0) and 1% (v/v) 1-butanol. The reaction was performed at 50°C, with shaking at 80 rpm in a small shaking incubator (JEIO TECH, Republic of Korea). Samples were measured in an external equilibrium solution, and the amount of product was quantitatively evaluated using a spectrophotometer.

### Analytical methods

Samples were analyzed using a C18 column (SunFireTM C18, 5 μm, 4.6 × 250 mm). Mobile phase A was distilled water containing 0.1% (v/v) formic acid (HPLC grade), and mobile phase B was 20% (v/v) acetonitrile in distilled water (HPLC grade). The mobile phase gradient was increased from 60 % to 80 % buffer B over 5 min, and then 100% buffer B was flowed for 10 min. The flow rate was set at 1 ml∙min^−1^. Products were detected using a UV/Vis detector (SPD-20A) at 260 nm and analyzed using the CMB-20A system controller (Shimadzu, Kyoto, Japan). Calibrations were conducted using 10 levels of rTA (Sigma-Aldrich, Missouri, USA) and MHET (AmBeed, Chicago, USA) in 50 mM glycine-NaOH (pH 8.7) and BHET (Sigma-Aldrich, Missouri, USA) in DMSO for blocking self-hydrolysis.

### Spectrophotometric assay

Quantitative determination of rTA was performed using a UV/VIS spectrophotometer (UV-1800, Shimadzu, Japan). All samples mixed with rTA, MHET, and BHET and then diluted to 10% (v/v) with 50 mM Glycine-NaOH (pH 8.7) solution. After adding *Is*MHETase^32^ (8 nM), the reaction mixtures were incubated at 37°C for 3 h or more. For measurement, each sample was loaded into a quartz cuvette of the UV spectrophotometer (path length 10 mm, Starna Scientific, London, England) and measured by scanning at a wavelength range of 250–360 nm. The value obtained at the baseline wavelength (360 nm) was subtracted from that obtained at the measurement wavelengths (250, 260, and 270 nm), and the resulting value was input into each calibration equation. Calibration was performed by diluting rTA in 50 mM glycine-NaOH (pH 8.7) and measuring at eight levels at 250 nm (0.2–2 μM rTA), 260 nm (0.2–2 μM rTA), and 270 nm (0.2–2 μM rTA) (Supplementary Fig. 12).

### Structure determination

Purified *Is*PETase variant proteins were further purified for crystallization via fast-protein liquid chromatography (FPLC). Eluted proteins were exchanged from buffer B into buffer A and concentrated at a high concentration (>1.0 mM) using a 10k concentration column (Amicon® Ultra-15 Centrifugal Filter Unit). Crystallization was achieved using the sitting-drop vapor diffusion method on an MRC crystallization plate (Molecular Dimensions). Each 1.0-μl drop of the protein solution was mixed with 1.0 μl of commercially available sparse matrix screens, such as Index, PEG ion, I and II (Hampton Research), and Wizard Classic, on the plate and then equilibrated with 50 μL of reservoir solution at 20°C. The best-quality crystals of the Is-4pC, Is-4pC+A180V, Is-4pC+P181V, Is-8p, Z1-PETase, and Z2-PETase variants were found in G4 of Wizard Classic, G4 of Index, A1 of PEG ion I, F7 of Index, H7 of PEG ion II, and A11 of PEG ion I, respectively. The highest-quality crystals were transferred to a cryo-protectant solution consisting of the well solution with 20% (v/v) glycerol. Crystals were scintillation-frozen in a nitrogen gas flow at 4°C to obtain X-ray diffraction data at an angle of 1°∙s^−1^ and wavelength of 0.97934 Å. Data were collected at the 7A beamline of the Pohang Accelerator Laboratory (PAL, Pohang, Republic of Korea) using a Quantum 270 CCD detector (ADSC, USA). All X-ray diffraction data were indexed, integrated, and scaled using the HKL2000 software package.^28^ Molecular replacement of the collected data for all variants was conducted using the CCP4^29^ version of MOLREP,^30^ with the deposited *Is*PETase structure (PDB code 5XJH) used as a search model. Further model building to align PDB files with the Fo-Fc map was performed manually using WinCoot^29^, and refinement was performed using CCP4 refmac5.^29^ All data statistics are summarized in Table S3. The final refined models of Is-4pC, Is-4pC+A180V, Is-4pC+P181V, Is-8p, Z1-PETase, and Z2-PETase were deposited in the Protein Data Bank with PDB codes of 8H5M, 8H5O, 8H5J, 8H5K, and 8H5L, respectively.^31^

### Heat inactivation experiment

Protein heat inactivation was investigated using a *p-*NPB hydrolysis assay. When *p-* NPB is hydrolyzed, butyrate and *p*-nitrophenolate ions are produced, and the latter can be measured via UV/VIS spectrophotometry at 409 nm. *Is*PETase variants (200 nM *Is*PETase mutant in 200 μl of 50 mM glycine-NaOH [pH 9.0]) were inactivated at 60°C or 70°C (1–36 h) and then chilled in an ice bath for 10 min. The reaction mixture contained 1% (v/v) Triton X-100, 10% MeOH, 50 mM phosphate buffered saline buffer (pH 7.8; Sigma-Aldrich, Missouri, USA), and 1 mM *p-*NPB. *Is*PETase mutants were added to the reaction mixture to initiate reaction. The protein concentration in the reaction mixture was 500 nM, and the final volume was adjusted to 500 μL with distilled water. The reaction was monitored for 60 s, and 10 s of the interval between linear-increasing sections were used as data. Each data point was calculated except for the baseline, which was obtained by measuring the self-hydrolysis of *p-* NPB. All experiments were performed in triplicate.

### Bioreactor system

A lab-scale bioreactor (MARDO-PDA, BIOCNS, Republic of Korea) was customized to analyze PET depolymerization. Three sets of jars with a capacity of 500 ml were deposited into the reactor. A non-contact type temperature sensor was used, and the reactor operated at a range of 30°C to 80°C. The temperature was regulated using a refrigeration and heating bath circulator (RW3-1025, JEIO TECH, Daejeon, Republic of Korea). The pH of the reaction solution was measured using a pH electrode (one-gel pH electrode with a digital display in 0.01 increments). The pH meter was calibrated at two pH levels (4.01 and 7.00). For the reaction, 2.5 μM of Z1-PETase was added to 150 ml of 10 mM Na_2_HPO_4_-HCl (pH 8.0) in the reactor. Samples of transparent and colored post-consumer PET powder were added at a concentration of 2.5% (w/v), and the sample and enzyme were mixed with an agitation speed of 200 rpm. The reaction solution was maintained at pH 8.0 through buffer titration with 0.3575 N NaOH (40%; extra pure grade, Duksan, Republic of Korea). The weight of added NaOH was measured using a precision weighting measurement system and a micro-precision scale (EK-410i; AND, Seoul, Republic of Korea). Samples were taken at specific time points and then filtered using a 0.22-μm PVDF syringe filter (SFPVDF013022CB, Futecs, Korea) to remove the residual PET powder. The obtained samples were diluted with 50 mM Glycine-NaOH (pH 9.0) and rTA, MHET, and BHET were detected using HPLC. The total amount of the released product was calculated, accounting for the amount of 0.3575 N NaOH added during the process.

## Supporting information

Supplementary Informations

## Acknowledgements

This research was supported by the Bio & Medical Technology Development Program of the National Research Foundation (NRF) funded by the Ministry of Science & ICT (NRF-2020M3A9I5037635) and the Cooperative Research Program for Agricultural Science & Technology Development (Project No. PJ01492602), Rural Development Administration, Republic of Korea. This research was also supported by the Technology Innovation Program (20018132, Development of the biodegradable polybutylene plastics, PBAT and PBS, from biomass) funded By the Ministry of Trade, Industry & Energy(MOTIE, Korea).

## Author contributions

K.-J. Kim, S.H. Lee, H. Seo, H. conceived the experiments. S.H. Lee, H. Seo, H. Hong, J. Park, D. Ki, and M. Kim performed PET hydrolysis experiments. S.H. Lee, H. Seo performed the structural determination. K.-J. Kim, S.H. Lee, Seo, H, and H-J Kim analyzed the data and wrote the paper. K.-J. Kim supervised the project.

## Reporting summery

Further information on research design is available in the Nature Research Reporting Summary linked to this article.

## Data availability

Data supporting the findings of this work are available within the paper and its Supplementary Information file. The methods are in the Supplementary information. The structures of *Is*PETase variants are deposited in Protein Data Bank with an accession code of 8H5M, 8H5N, 8H5O, 8H5J, 8H5K, and 8H5L. The source data underlying Figs. 1b, 1c, 2, 4 and 5 are associated with raw data. The datasets generated and analyzed during the current study are available from the corresponding authors upon request.

