## Supplementary Informations for "Balance-directed protein engineering of *Is*PETase enhances both PET hydrolysis activity and thermostability"

This PDF file includes: Supplementary Figures 1 to 12.
Supplementary Tables 1 to 3

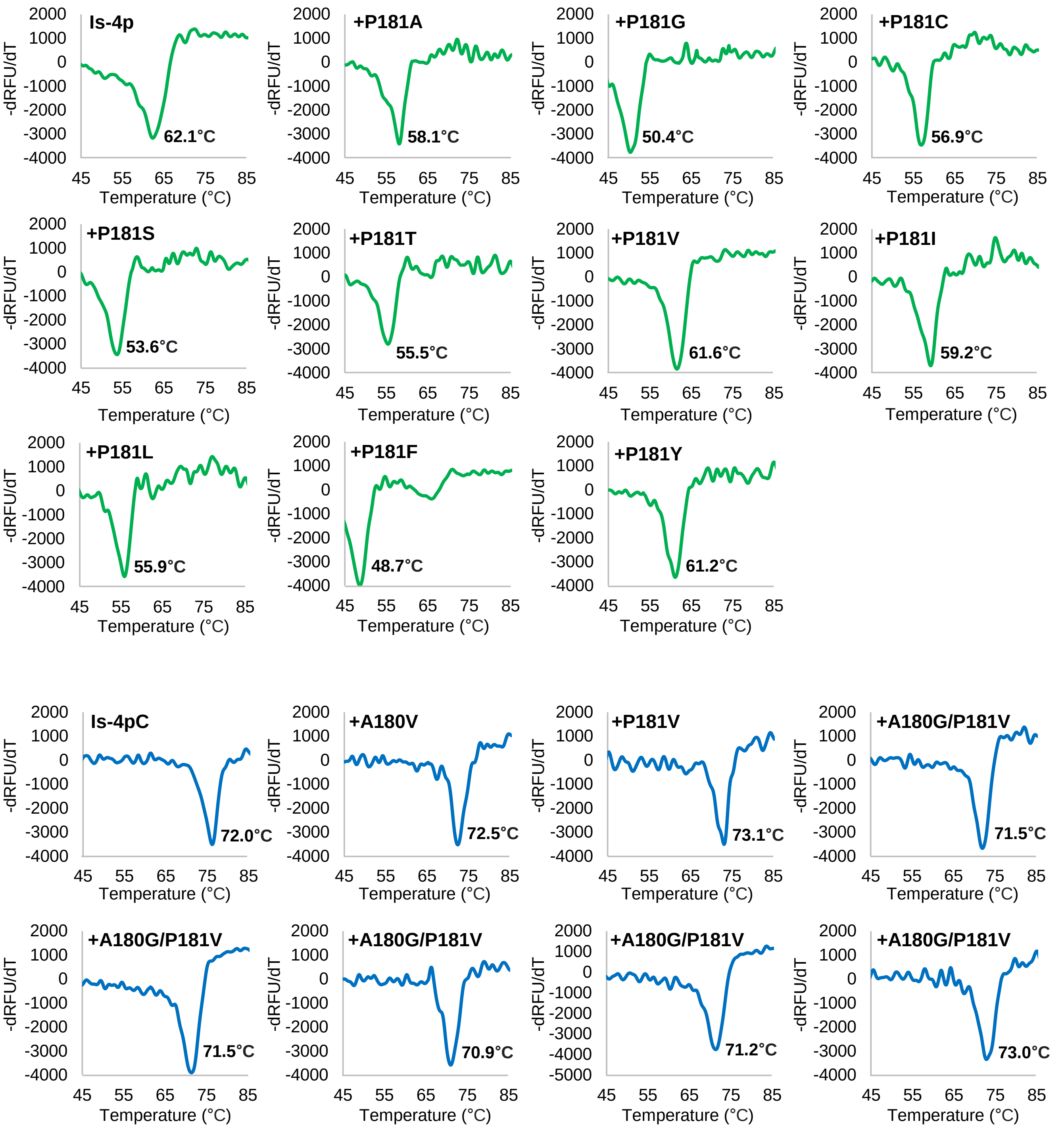


**Supplementary Figure 1. Differential scanning fluorimetry of the *Is*PETase variants used in this study.**

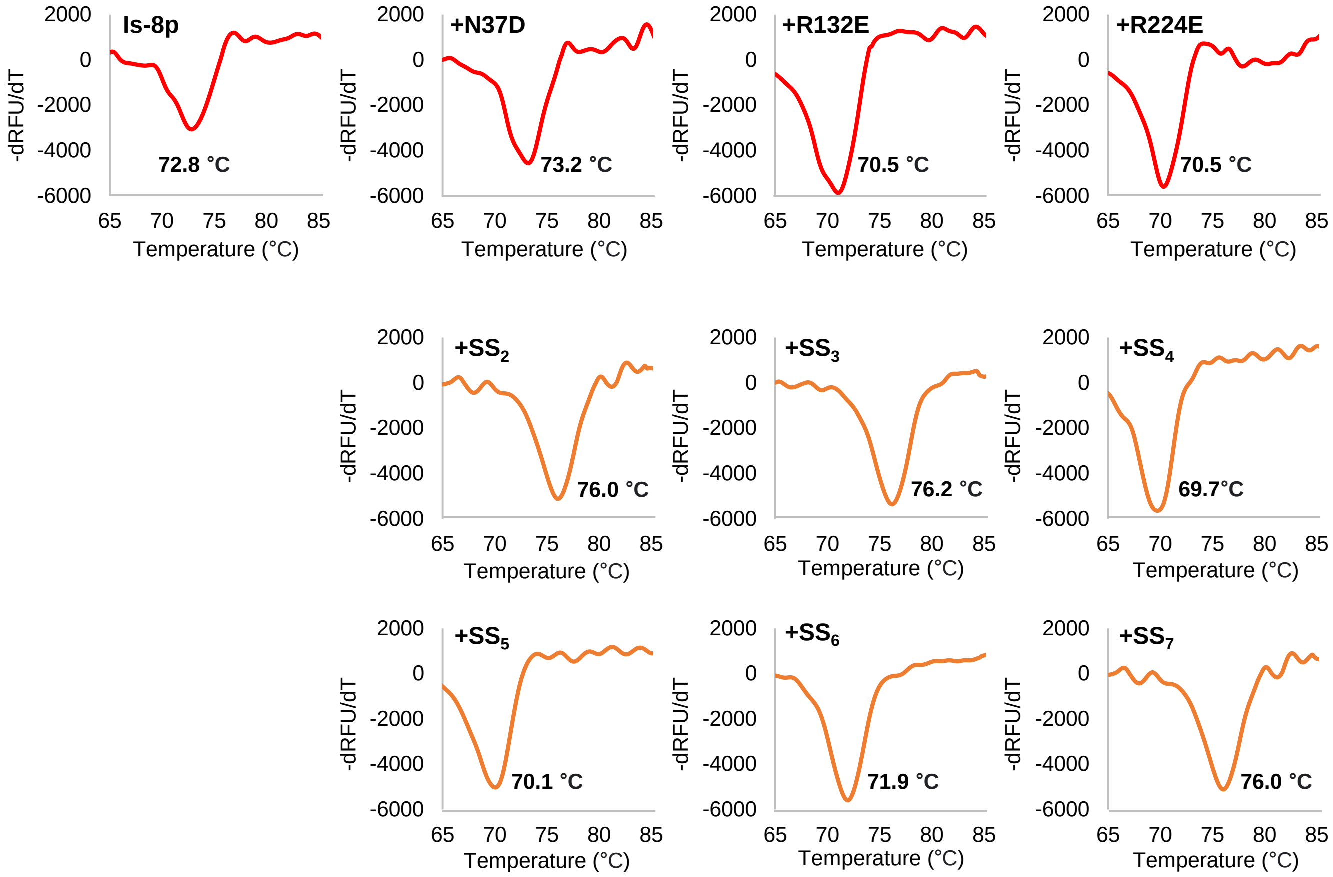


**Supplementary Figure 1. Differential scanning fluorimetry of the *Is*PETase variants used in this study. (continued)**

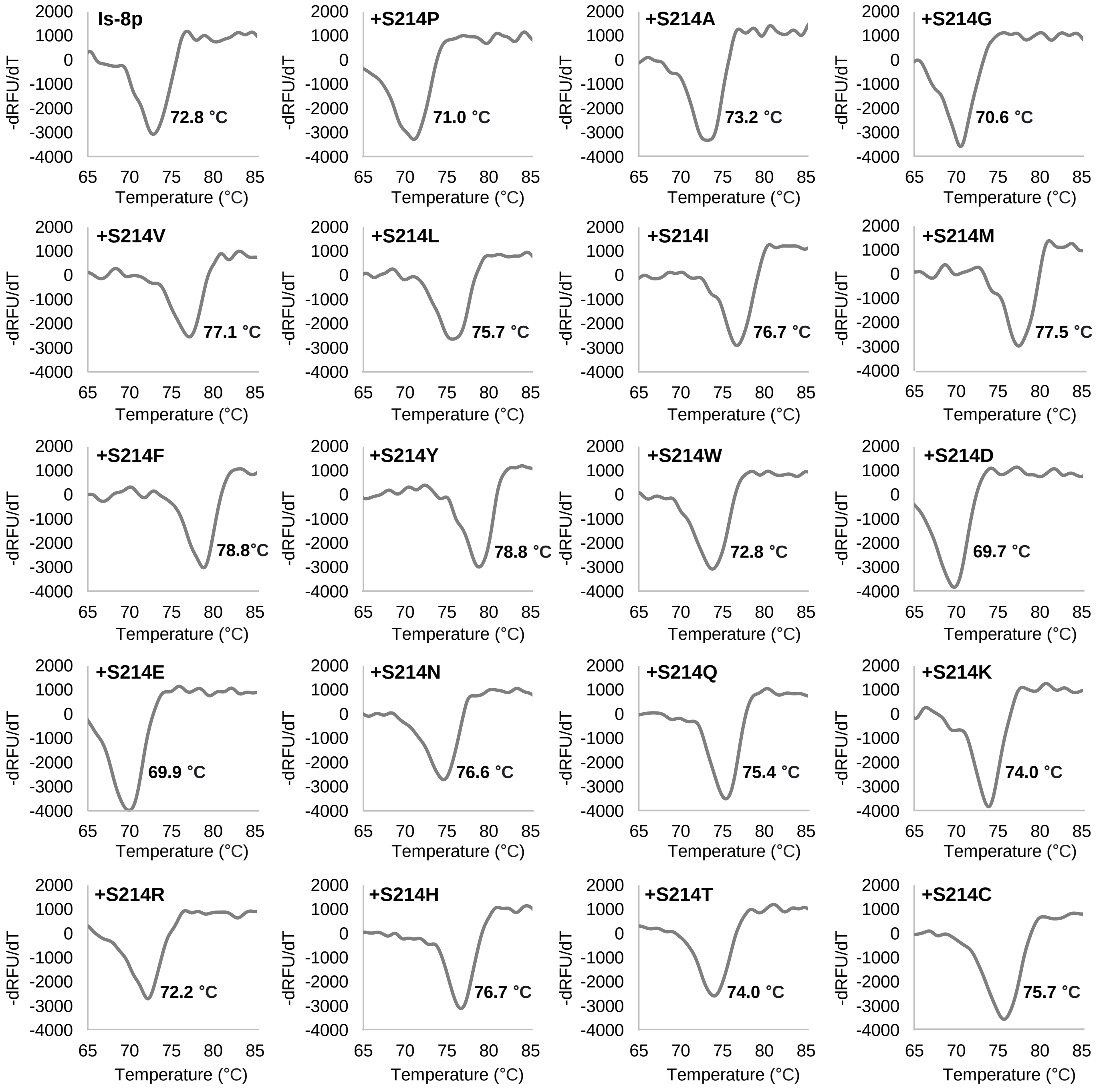


**Supplementary Figure 1. Differential scanning fluorimetry of the *Is*PETase variants used in this study. (continued)**

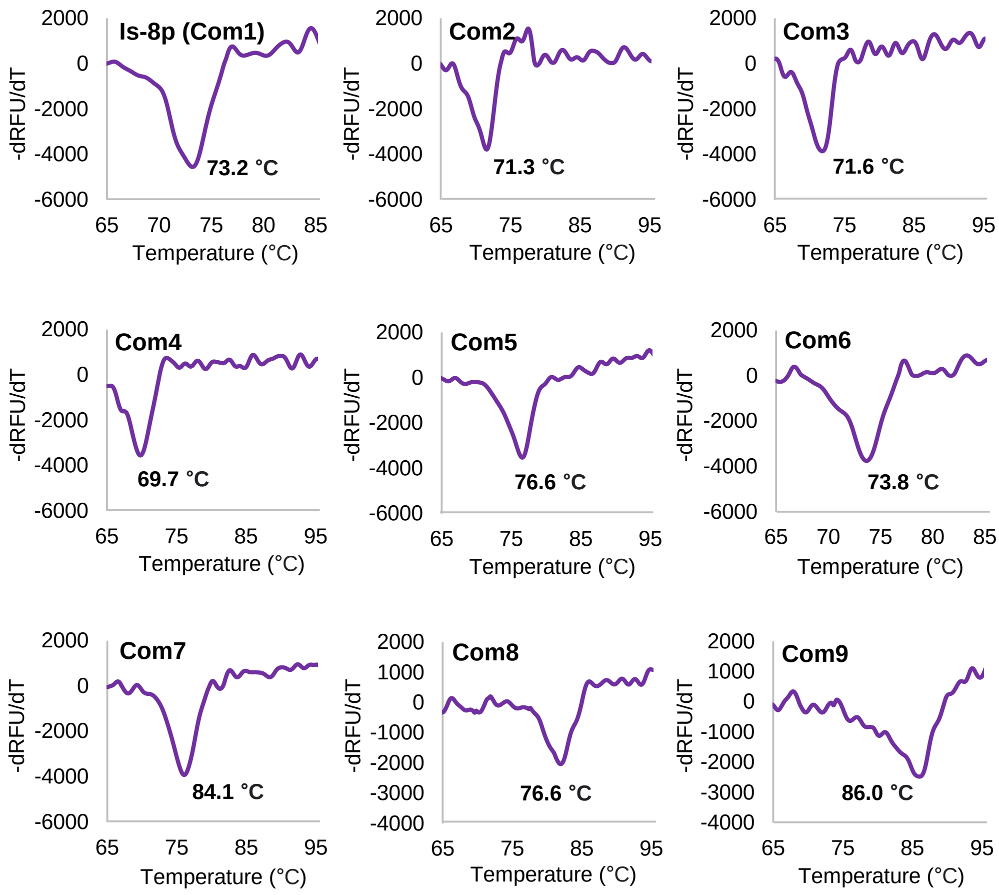


**Supplementary Figure 1. Differential scanning fluorimetry of the *Is*PETase variants used in this study. (continued)**

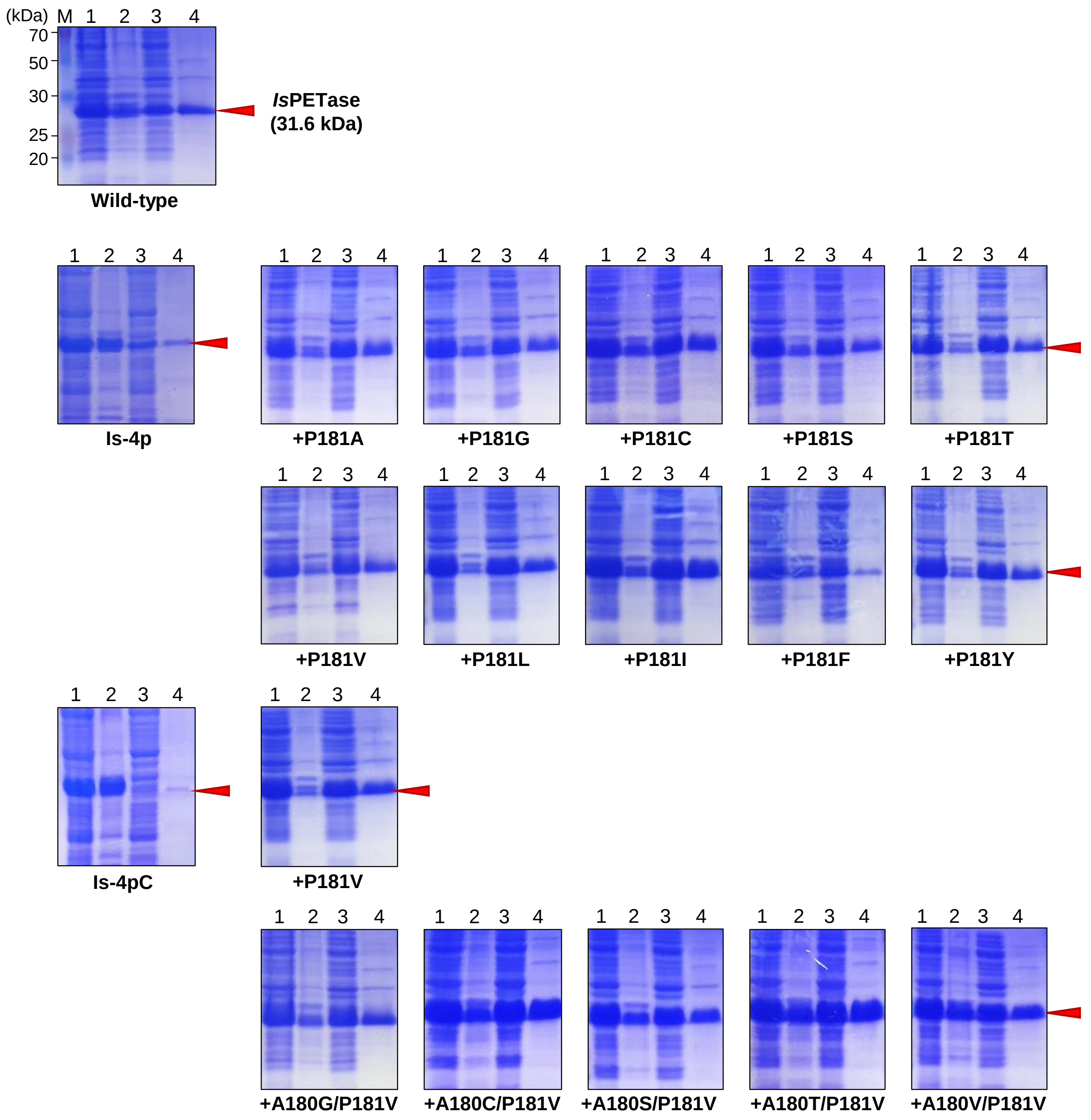


**Supplementary Figure 2. SDS-page analysis of overexpression of Is-4p+P181X and Is-4pC+A180X/P181V variants.** The purified *Is*PETase variants using the *E. coli* Rosetta-gami™ 2(DE3) expression system are loaded on SDS-PAGE. The molecular weight of the protein is 31.6 kDa. Lane M represents the ladder (molecular marker). Lanes 1, 2, 3, and 4 indicate cell lysate, cell pellets, and supernatant, and eluted protein, respectively.

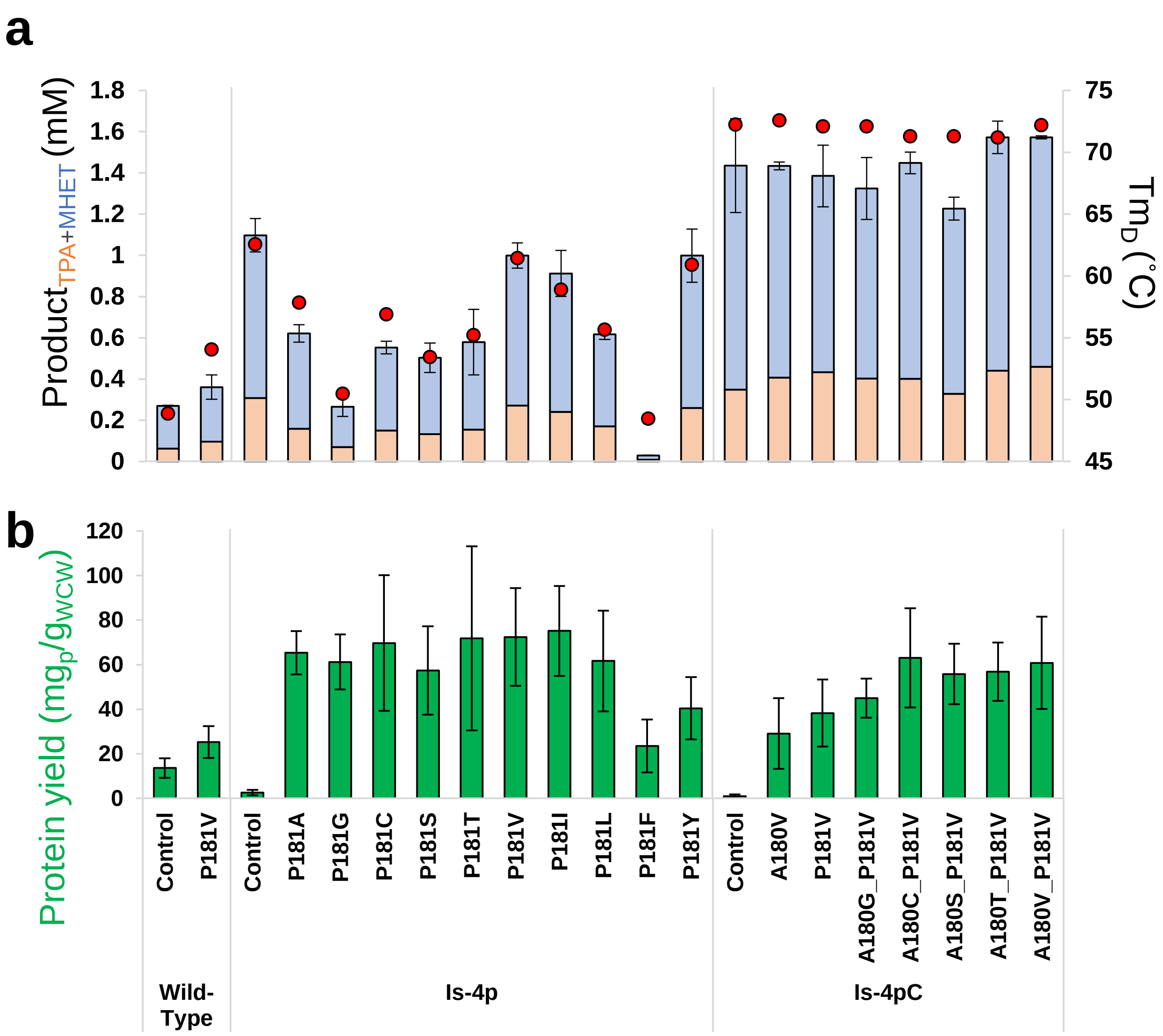


**Supplementary Figure 3. PET degradation activity, thermal stability ($Tm_D$), and protein yield of 4p+P181X and 4pC+A180X/P181V variants. a,** PET hydrolysis activity and thermal stability of Is-4p+P181X and Is-4pC+A180X+P181V. The PET hydrolysis activity is measured at 6 hours of incubation using 15 $gL^{-1}$ of bottle-derivative PET powder in 50 mM glycine-NaOH pH 9.0 at 50 °C. All the data is reacted in triplicate via HPLC. **b,** Protein yields of Is-4p+P181X and Is-4pC+A180X+P181V. The protein yields are detected from soluble protein expression in the elution buffer. The unit is protein weight ($mg_p$) of 1g of wet cell weight ($g_{wcw}$). All the experiments were performed in triplicate, and the error bars of measuring samples represent the sampling s.d. (n=3).

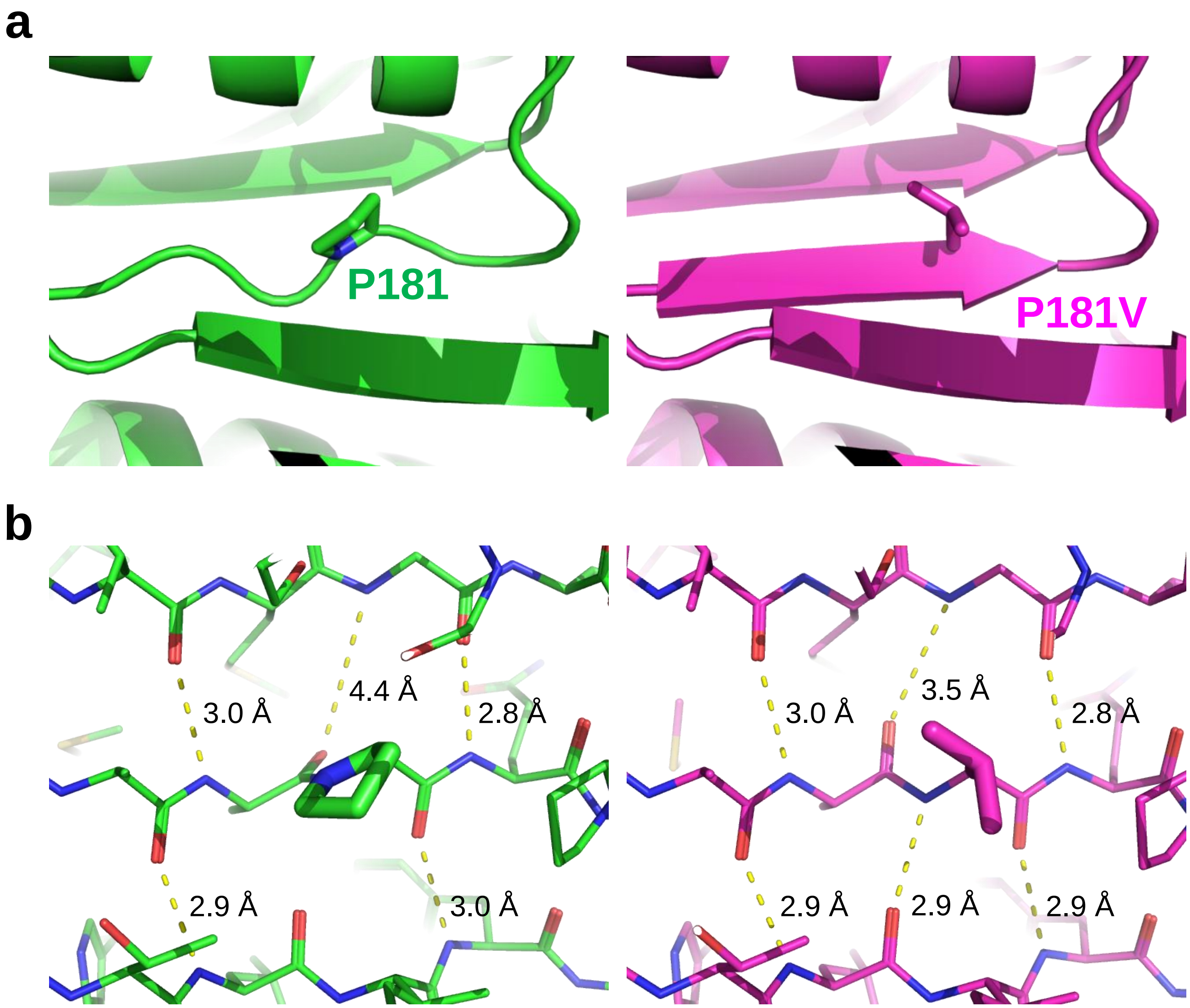


**Supplementary Figure 4. β-strand reconstruction in the *Is*PETase+P181V variant. a,** Comparison of β-strand of Is-4pC and Is-4pC+P181V. The structures of Is-4pC and Is-4pC+P181V are colored as green and magenta, respectively. The P181 and P181A residue are shown as a stick. **b,** Comparison of hydrogen bond length. The hydrogen bonds are shown as a yellow-colored dotted- lines and their length are labeled.

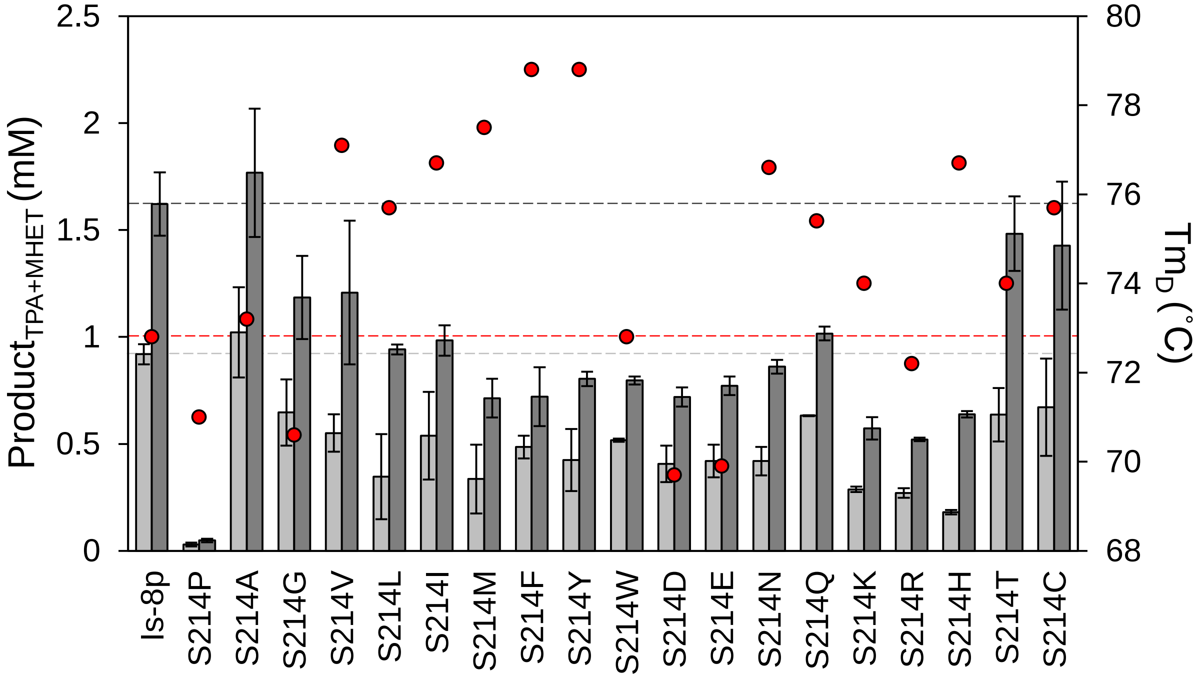


**Supplementary Figure 5. Site saturation mutagenesis of Ser214.** The PET hydrolysis activity is measured at 3 and 6 hours of incubation using bottle-derivative PET at 50 °C colored as light gray and gray, respectively. All experiments were performed in triplicate and detected via HPLC, and error bars of measuring samples represent the sampling s.d. (n=3).

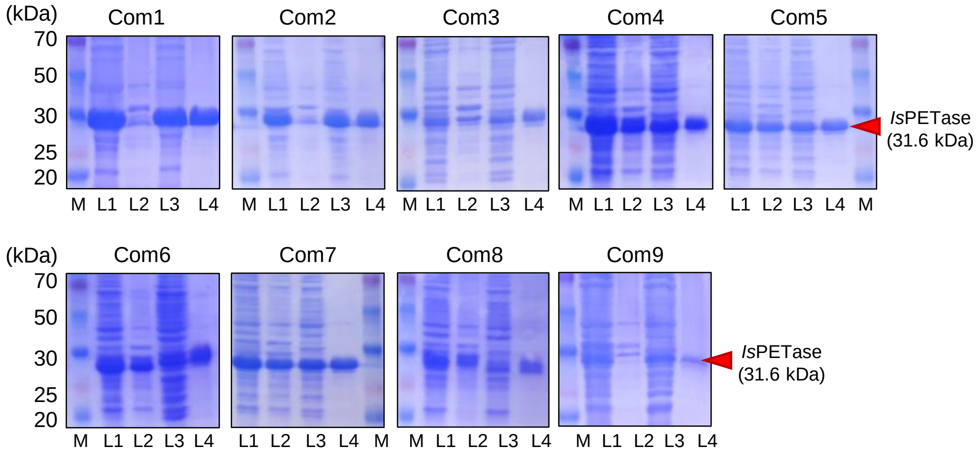


**Supplementary Figure 6. Analysis of overexpression of combinatorial variants on the 8p variants.** The purified *Is*PETase combinatorial mutants using the *E. coli* Rosetta-gami™ 2(DE3) expression system are loaded on SDS-PAGE. The molecular weight of the protein is 31.6 kDa. Lane M represents the ladder (molecular marker). Lanes 1, 2, 3, and 4 are cell lysate, cell pellets, supernatant, and eluted protein, respectively.

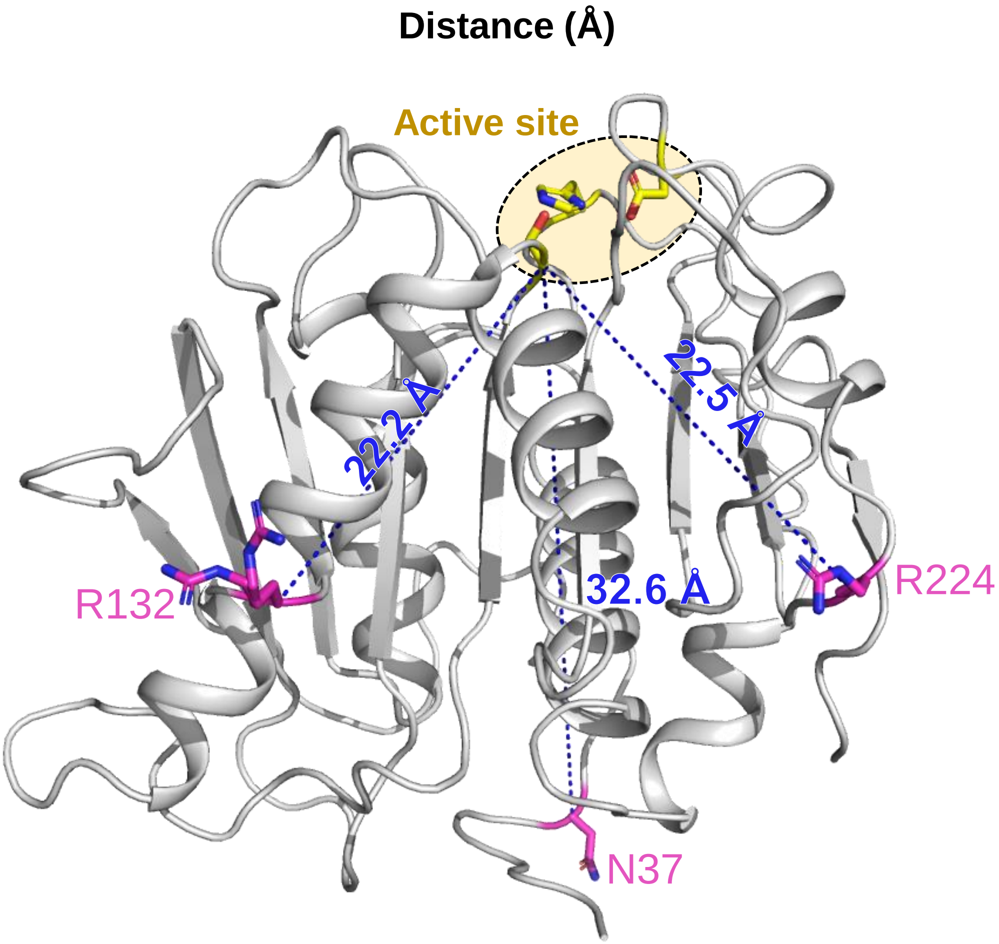


**Supplementary Figure 7. The distance between the catalytic site and the residues of N37D, R132E, and R224E.** The Is-4pC variant structure is shown as a cartoon diagram colored gray. The catalytic and surface charge mutation residues are shown as a stick with yellow and magenta colores, respectively.

a

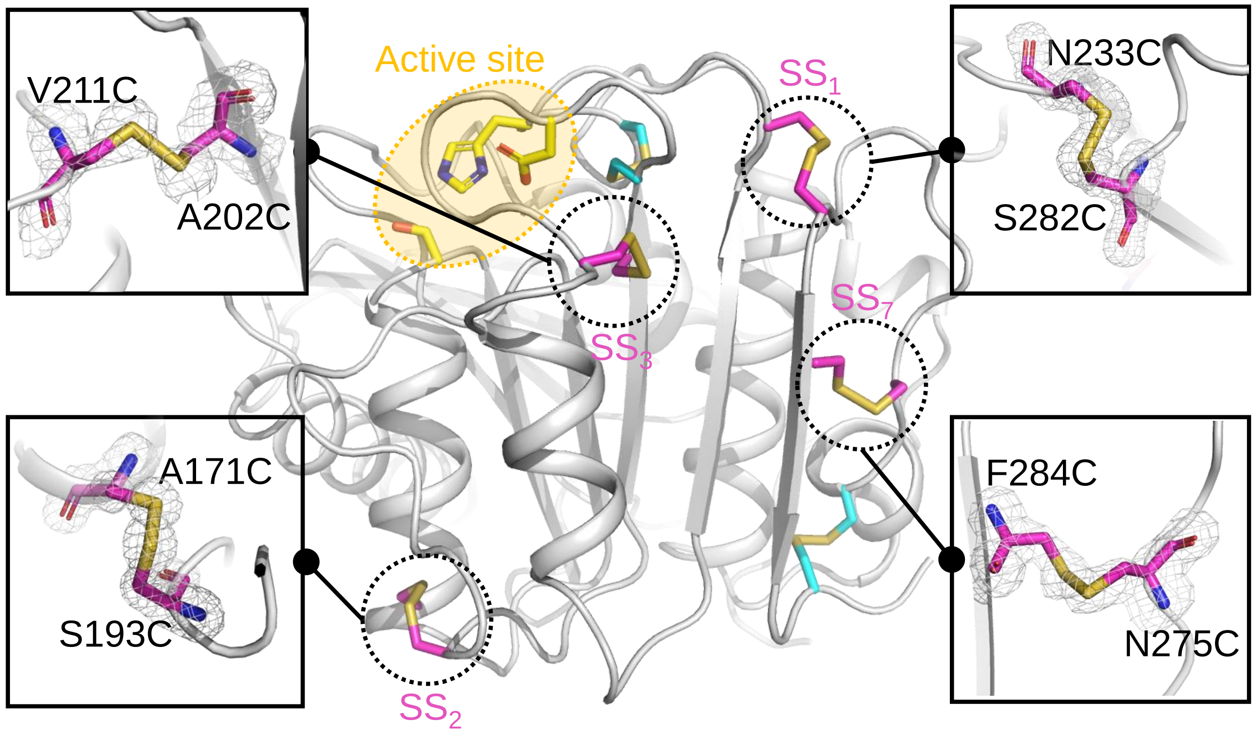


b

| Pair of Cysteine | | Chi1(X1) | Chi2(X2) | Chi2'(X2') | Chi1'(X1') | $C^{\beta}$–$S^{\gamma}$–$S^{\gamma}$–$C^{\beta}$ (°) | Distance (Å) | Disulfide Strain Energy (kJ/mol) |
|---|---|---|---|---|---|---|---|---|
| A171C | S193C | -70.88 (±1.67) | -129.36 (±2.85) | 86.32 (±1.12) | 53.20 (±1.07) | 66.34 (±0.87) | 2.12 (±0.04) | 17.95 (±0.61) |
| A202C | V211C | 77.47 (±1.93) | 105.96 (±2.87) | -90.56 (±4.02) | -65.52 (±2.92) | 111.37 (±1.30) | 2.30 (±0.02) | 23.91 (±0.58) |
| N233C | S282C | -61.63 (±3.30) | -55.92 (±3.21) | 62.20 (±4.09) | -91.95 (±6.90) | 110.60 (±2.53) | 2.06 (±0.03) | 17.88 (±2.53) |
| N275C | F284C | -74.05 (±0.31) | -62.53 (±3.21) | -115.12 (±2.32) | -178.83 (±0.90) | -79.89 (±1.60) | 2.10 (±0.02) | 12.63 (±0.31) |

**Supplementary Figure 8. Disulfide bond formation analysis of Z2-PETase.** The Z2-PETase structure is shown as a cartoon diagram colored gray. Disulfide bonds from the *Is*PETase$^{Wild\text{-}type}$ and constructed in this study are shown as a stick with cyan and magenta colored, respectively. The electron density (Fo-Fc) maps of the introduced disulfide bonds are shown with a 1.5 σ contour.

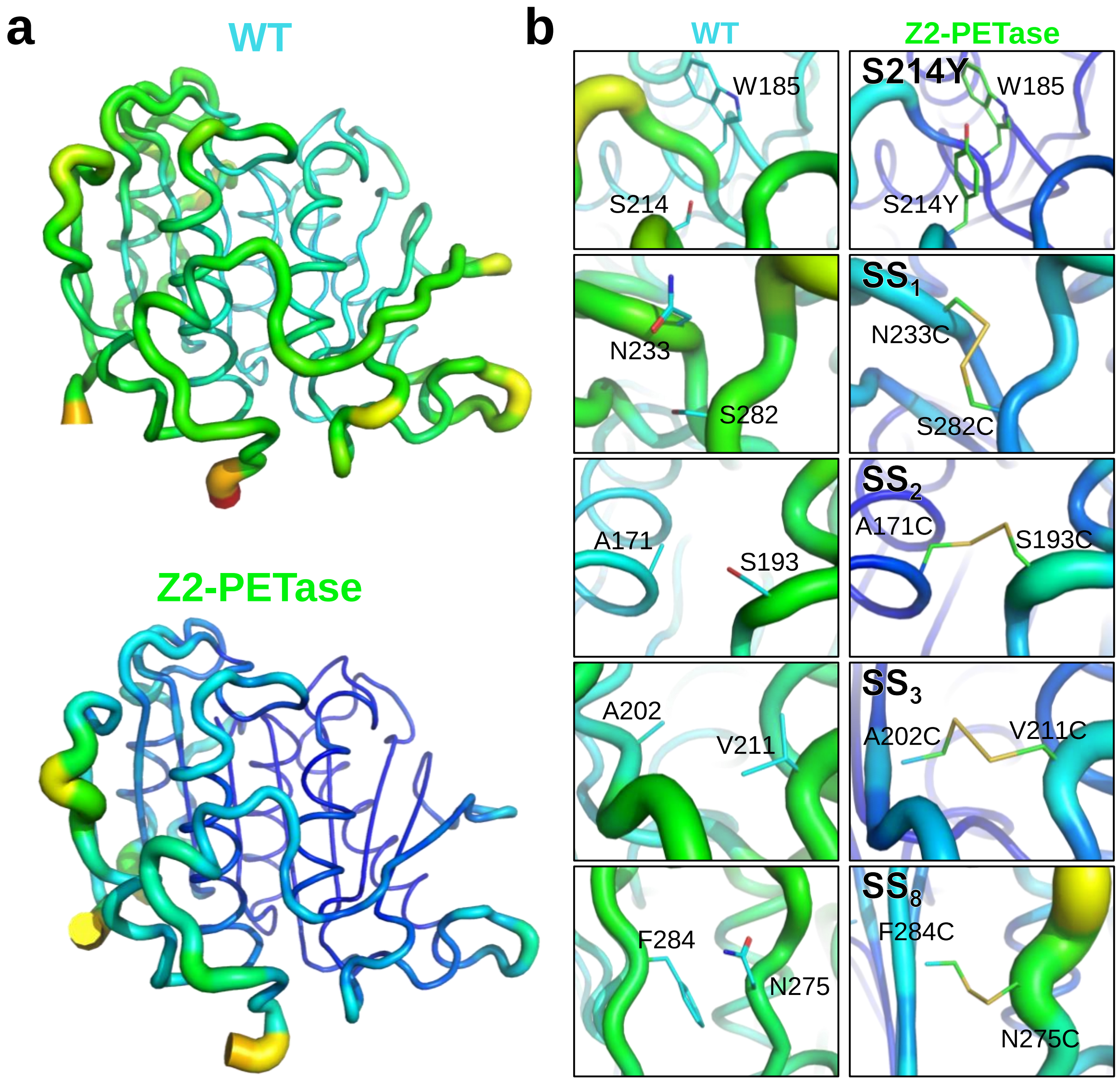


**Supplementary Figure 9. Comparison of b-factor putty of *Is*PETase$^{\text{Wild-type}}$ and Z2-PETase.**

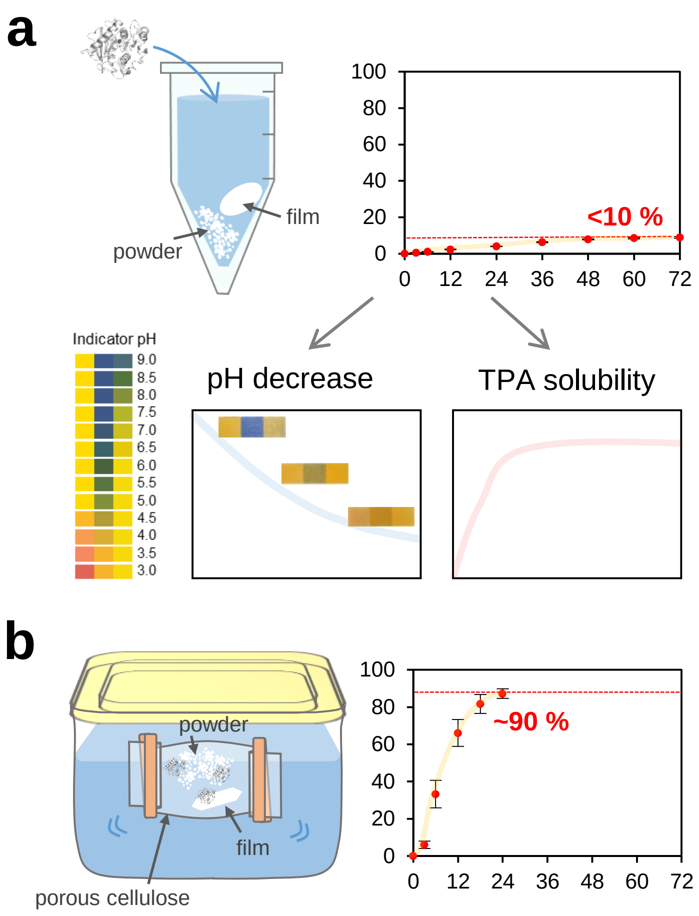


**Supplementary Figure 10. Schematic diagram of a product-deficient dialysis system. a,** The limitation to measuring real PET hydrolysis activity at the closed vial system. **b,** Designed product deficient *in-situ* dialysis system.

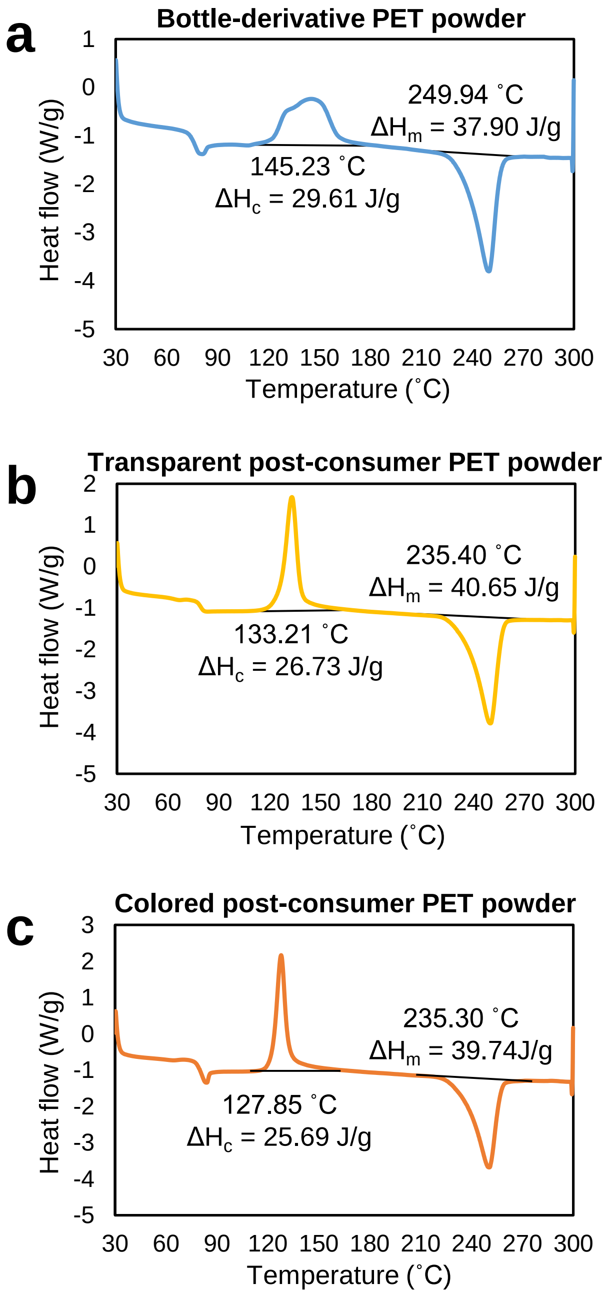


**Supplementary Figure 11. A DSC plot for polyethylene terephthalate (PET) sample.** The crystallinity of bottle-derivative PET, Transparent post-consumer PET, and Colored post-consumer PET powder were calculated using DSC experiments. The crystallinity of bottle-derivative PET, Transparent post-consumer PET, and Colored post-consumer PET powder were calculated using DSC experiments. The bottle-derivative PET has an enthalpy of cold crystallization ($\Delta H_c$) of 29.61 J/g, a melting enthalpy ($\Delta H_m$) of 37.90 J/g, and a percent crystallinity of 7.8 %. The Transparent post-consumer PET has an enthalpy of cold crystallization ($\Delta H_c$) of 26.73 J/g, a melting enthalpy ($\Delta H_m$) of 40.65 J/g, and a percent crystallinity of 10.1 %. The colored post-consumer PET powder has an enthalpy of cold crystallization ($\Delta H_c$) of 25.69 J/g, a melting enthalpy ($\Delta H_m$) of 39.74 J/g, and a percent crystallinity of 10.04 %.

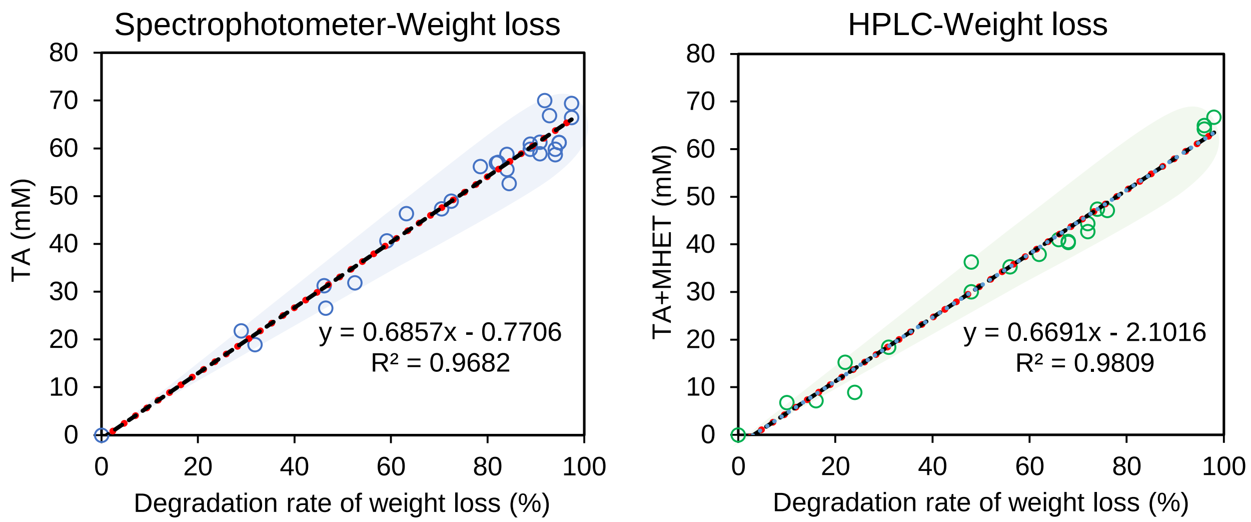


**Supplementary Figure 12. PET hydrolysis rate calculation comparison between HPLC and spectrophotometer detection method with weight loss of the PET powder.** The total amount of each sample was starting from 50 mg. The x-slope of the spectrophotometer and the HPLC is within error range.

#### Supplementary Table 1. List of primers used in this study.

| Site-directed mutagenesis | Forward sequence (5’-3’) | Reverse sequence (5’-3’) |
|---|---|---|
| N37D | CATATGCGCGGCCCGGATCCGACAGCCGCCAGT | ACTGGCGGCTGTCGGATCCGGGCCGCGCATATG |
| R132E | CAAATGGCCGCGCTGGAACAGGTGGCGTCGTTA | TAACGACGCCACCTGTTCCAGCGCGGCCATTTG |
| A171C | TCCCTGATCTCTGCTTGCAACAACCCTTCGCTG | CAGCGAAGGGTTGTTGCAAGCAGAGATCAGGGA |
| A180V | TCGCTGAAAGCAGCGGTGCCTCAAGCACCATGG | CCATGGTGCTTGAGGCACCGCTGCTTTCAGCGA |
| A180G+P181V | TCGCTGAAAGCAGCGGCGGTGCAAGCACCATGG | TCGCTGAAAGCAGCGGTGCCTCAAGCACCATGG |
| A180C+P181V | TCGCTGAAAGCAGCGTGCGTGCAAGCACCATGG | TCGCTGAAAGCAGCGGTGCCTCAAGCACCATGG |
| A180S+P181V | TCGCTGAAAGCAGCGAGCGTGCAAGCACCATGG | TCGCTGAAAGCAGCGGTGCCTCAAGCACCATGG |
| A180T+P181V | TCGCTGAAAGCAGCGACCGTGCAAGCACCATGG | TCGCTGAAAGCAGCGGTGCCTCAAGCACCATGG |
| A180V+P181V | TCGCTGAAAGCAGCGGTGGTGCAAGCACCATGG | TCGCTGAAAGCAGCGGTGCCTCAAGCACCATGG |
| P181A | CTGAAAGCAGCGGCGGCGCAAGCACCATGGGAT | ATCCCATGGTGCTTGGACCGCCGCTGCTTTCAG |
| P181G | CTGAAAGCAGCGGCGGGCCAAGCACCATGGGAT | ATCCCATGGTGCTTGGACCGCCGCTGCTTTCAG |
| P181C | CTGAAAGCAGCGGCGTGCCAAGCACCATGGGAT | ATCCCATGGTGCTTGGACCGCCGCTGCTTTCAG |
| P181S | CTGAAAGCAGCGGCGAGCCAAGCACCATGGGAT | ATCCCATGGTGCTTGGACCGCCGCTGCTTTCAG |
| P181T | CTGAAAGCAGCGGCGACCCAAGCACCATGGGAT | ATCCCATGGTGCTTGGACCGCCGCTGCTTTCAG |
| P181V | CTGAAAGCAGCGGCGGTCCAAGCACCATGGGAT | ATCCCATGGTGCTTGGACCGCCGCTGCTTTCAG |
| P181I | CTGAAAGCAGCGGCGATTCAAGCACCATGGGAT | ATCCCATGGTGCTTGGACCGCCGCTGCTTTCAG |
| P181L | CTGAAAGCAGCGGCGCTGCAAGCACCATGGGAT | ATCCCATGGTGCTTGGACCGCCGCTGCTTTCAG |
| P181F | CTGAAAGCAGCGGCGTTTCAAGCACCATGGGAT | ATCCCATGGTGCTTGGACCGCCGCTGCTTTCAG |
| P181Y | CTGAAAGCAGCGGCGTATCAAGCACCATGGGAT | ATCCCATGGTGCTTGGACCGCCGCTGCTTTCAG |
| S193C | TCGACAAATTTTAGTTGCGTAACTGTGCCCACG | CGTGGGCACAGTTACGCAACTAAAATTTGTCGA |
| A202C | CCCACGCTGATCTTCTGCTGTGAAAACGATAGT | ACTATCGTTTTCACAGCAGAAGATCAGCGTGGG |
| V211C | GATAGTATAGCCCCGTGCAACTCTTCAGCACTT | AAGTGCTGAAGAGTTGCACGGGGCTATACTATC |
| S214Y | GCCCCGGTCAACTCTTATGCACTTCCTATCTAT | ATAGATAGGAAGTGCATAAGAGTTGACCGGGGC |
| R224E | TATGATTCTATGTCAGAAAACGCTAAGCAGTTT | AAACTGCTTAGCGTTTTCTGACATAGAATCATA |
| N233C | CAGTTTCTCGAAATTTGCGGTGGCTCACATTCC | GGAATGTGAGCCACCGCAAATTTCGAGAAACTG |
| N275C | ACTTTTGCCTGCGAGTGCCCGAATAGCACCAGA | TCTGGTGCTATTCGGGCACTCGCAGGCAAAAGT |
| S282C | AATAGCACCAGAGTGTGCGATTTTCGTACAGCG | CGCTGTACGAAAATCGCACACTCTGGTGCTATT |
| F284C | ACCAGAGTGAGCGATTGCCGTACAGCGAATTGC | GCAATTCGCTGTACGGCAATCGCTCACTCTGGT |
| S282C+F284C | ACCAGAGTGTGCGATTGCCGTACAGCGAATTGC | GCAATTCGCTGTACGGCAATCGCACACTCTGGT |

**Supplementary Table 2. DNA sequence of the *Is*PETase variant.**

| Enzyme | Nucleotide sequence |
|---|---|
| 4p-PETase | ATGCGCGGTCCGAATCCGACAGCCGCCAGTTTGGAAGCGAGCGCTGGTCCATTCACCGTTCGCTCC<br>TTTACCGTGAGTAGACCGAGCGGTTATGGCGCTGGCACCGTTTACTATCCAACAAATGCTGGGGGT<br>ACCGTGGGCGCCATAGCCATAGTTCCCGGGTATACGGCACGGCAGTCATCAATTAAATGGTGGGGA<br>CCGCGTCTGGCATCCCACGGTTTCGTAGTAATTACAATTGACACAAATTCCACGTTAGACCAGCCA<br>GAAAGTCGGAGTTCGCAACAAATGGCCGCGCTGCGCCAGGTGGCGTCGTTAAACGGTACAAGTAG<br>CAGCCCGATTTACGGAAAGGTCGATACCGCTCGTATGGGTGTTATGGGGTGGAGTATGGGAGGTGG<br>AGGCTCCCTGATCTCTGCTGCTAACAACCCTTCGCTGAAAGCAGCGGCGCCTCAAGCACCATGGC<br>ATTCTTCGACAAATTTTAGTTCTGTAACTGTGCCCACGCTGATCTTCGCATGTGAAAACGATAGTAT<br>AGCCCCGGTCAACTCTTCAGCACTTCCTATCTATGATTCTATGTCACGCAACGCTAAGCAGTTTCTC<br>GAAATTAACGGTGGCTCACATTCCTGTGCGAATACCGGCAATTCTGACCAAGCATTAATCGGAAAA<br>AAAGGCGTTGCATGGATGAAACGTTTTATGGACAATGATACTAGGTATTCTACTTTTGCCTGCGAGA<br>ACCCGAATAGCACCAGAGTGAGCGATTTTCGTACAGCGAATTGCAGC |
| 4pC-<br>PETase | ATGCGCGGTCCGAATCCGACAGCCGCCAGTTTGGAAGCGAGCGCTGGTCCATTCACCGTTCGCTCC<br>TTTACCGTGAGTAGACCGAGCGGTTATGGCGCTGGCACCGTTTACTATCCAACAAATGCTGGGGGT<br>ACCGTGGGCGCCATAGCCATAGTTCCCGGGTATACGGCACGGCAGTCATCAATTAAATGGTGGGGA<br>CCGCGTCTGGCATCCCACGGTTTCGTAGTAATTACAATTGACACAAATTCCACGTTAGACCAGCCA<br>GAAAGTCGGAGTTCGCAACAAATGGCCGCGCTGCGCCAGGTGGCGTCGTTAAACGGTACAAGTAG<br>CAGCCCGATTTACGGAAAGGTCGATACCGCTCGTATGGGTGTTATGGGGTGGAGTATGGGAGGTGG<br>AGGCTCCCTGATCTCTGCTGCTAACAACCCTTCGCTGAAAGCAGCGGCGCCTCAAGCACCATGGC<br>ATTCTTCGACAAATTTTAGTTCTGTAACTGTGCCCACGCTGATCTTCGCATGTGAAAACGATAGTAT<br>AGCCCCGGTCAACTCTTCAGCACTTCCTATCTATGATTCTATGTCACGCAACGCTAAGCAGTTTCTC<br>GAAATTTGCGGTGGCTCACATTCCTGTGCGAATACCGGCAATTCTGACCAAGCATTAATCGGAAAA<br>AAAGGCGTTGCATGGATGAAACGTTTTATGGACAATGATACTAGGTATTCTACTTTTGCCTGCGAGA<br>ACCCGAATAGCACCAGAGTGTGCGATTTTCGTACAGCGAATTGCAGC |
| Z1-PETase | ATGCGCGGCCCGGATCCGACAGCCGCCAGTTTGGAAGCGAGCGCTGGTCCATTCACCGTTCGCTC<br>CTTTACCGTGAGTAGACCGAGCGGTTATGGCGCTGGCACCGTTTACTATCCAACAAATGCTGGGGG<br>TACCGTGGGCGCCATAGCCATAGTTCCCGGGTATACGGCACGGCAGTCATCAATTAAATGGTGGGG<br>ACCGCGTCTGGCATCCCACGGTTTCGTAGTAATTACAATTGACACAAATTCCACGTTAGACCAGCC<br>AGAAAGTCGGAGTTCGCAACAAATGGCCGCGCTGGAACAGGTGGCGTCGTTAAACGGTACAAGT<br>AGCAGCCCGATTTACGGAAAGGTCGATACCGCTCGTATGGGTGTTATGGGGTGGAGTATGGGAGGT<br>GGAGGCTCCCTGATCTCTGCTTGCAACAACCCTTCGCTGAAAGCAGCGGTGGTGCAAGCACCATG<br>GCATTCTTCGACAAATTTTAGTTGCGTAACTGTGCCCACGCTGATCTTCGCATGTGAAAACGATAGT<br>ATAGCCCCGGTCAACTCTTCAGCACTTCCTATCTATGATTCTATGTCAGAAAACGCTAAGCAGTTTC<br>TCGAAATTTGCGGTGGCTCACATTCCTGTGCGAATACCGGCAATTCTGACCAAGCATTAATCGGAA<br>AAAAAGGCGTTGCATGGATGAAACGTTTTATGGACAATGATACTAGGTATTCTACTTTTGCCTGCGA<br>GAACCCGAATAGCACCAGAGTGTGCGATTTTCGTACAGCGAATTGCAGC |
| Z2-PETase | ATGCGCGGCCCGGATCCGACAGCCGCCAGTTTGGAAGCGAGCGCTGGTCCATTCACCGTTCGCTC<br>CTTTACCGTGAGTAGACCGAGCGGTTATGGCGCTGGCACCGTTTACTATCCAACAAATGCTGGGGG<br>TACCGTGGGCGCCATAGCCATAGTTCCCGGGTATACGGCACGGCAGTCATCAATTAAATGGTGGGG<br>ACCGCGTCTGGCATCCCACGGTTTCGTAGTAATTACAATTGACACAAATTCCACGTTAGACCAGCC<br>AGAAAGTCGGAGTTCGCAACAAATGGCCGCGCTGGAACAGGTGGCGTCGTTAAACGGTACAAGT<br>AGCAGCCCGATTTACGGAAAGGTCGATACCGCTCGTATGGGTGTTATGGGGTGGAGTATGGGAGGT<br>GGAGGCTCCCTGATCTCTGCTTGCAACAACCCTTCGCTGAAAGCAGCGGTGGTGCAAGCACCATG<br>GCATTCTTCGACAAATTTTAGTTGCGTAACTGTGCCCACGCTGATCTTCTGCTGTGAAAACGATAGT<br>ATAGCCCCGTGCAACTCTTATGCACTTCCTATCTATGATTCTATGTCAGAAAACGCTAAGCAGTTTCT<br>CGAAATTTGCGGTGGCTCACATTCCTGTGCGAATACCGGCAATTCTGACCAAGCATTAATCGGAAA<br>AAAAGGCGTTGCATGGATGAAACGTTTTATGGACAATGATACTAGGTATTCTACTTTTGCCTGCGAG<br>TGCCCGAATAGCACCAGAGTGTGCGATTGCCGTACAGCGAATTGCAGC |

### Supplementary Table 3. Data Collection and Refinement Statistics

| Protein | 4pC | 4pC+P181V | 8p-PETase | Z1-PETase | Z2-PETase |
|---|---|---|---|---|---|
| PDB deposit | 8H5M | 8H5O | 8H5J | 8H5K | 8H5L |
| Molecules | 1 | 1 | 2 | 1 | 3 |
| Space group | $P2_12_12_1$ | $P2_12_12_1$ | $C_12_1$ | $P2_12_12_1$ | $P12_11$ |
| **Unit cell** | | | | | |
| a, b, c (Å) | 50.7, 67.8, 78.5 | 50.9.52.6,95.2 | 139.2, 50.7, 80.9 | 51.0, 51.7, 65.6 | 100.2, 55.6, 100.4 |
| r (˚) | 90.00 | 90.00 | 90.0, 124.6, 90.0 | 90.00 | 90.0, 119.1, 90.0 |
| X-ray source | PAL-7A | PAL-7A | PAL-7A | PAL-7A | PAL-7A |
| Wavelength (Å) | 0.97934 | 0.97934 | 0.97934 | 0.97934 | 0.97934 |
| Energy (keV) | 12.660 | 12.660 | 12.660 | 12.660 | 12.660 |
| Resolution (Å) | 28.23-1.90 | 29.01-2.10 | 28.65-1.40 | 25.05-1.20 | 27.97-1.70 |
| **Reflections** | | | | | |
| Observed | 1213143 | 597332 | 828222 | 1391065 | 1181822 |
| Unique | 22366 | 15494 | 90308 | 63488 | 99271 |
| Completeness[1] | 94.6 (90.6) | 100.0 (99.8) | 98.3 (96.8) | 99.9 (99.2) | 97.1 (92.3) |
| $< I / \sigma(I) >$[1] | 22.79 (4.74) | 45.07 (16.61) | 30.00 (2.88) | 37.55 (2.78) | 18.55 (3.45) |
| $R_{merge}$ (%)[1] | 9.3 (67.6) | 11.4 (30.0) | 9.8 (51.3) | 6.4 (58.2) | 15.9 (80.0) |
| CC1/2[1] | 1.00 (0.62) | 0.99 (0.95) | 0.98 (0.74) | 1.00 (0.84) | 0.98 (0.73) |
| Protein atoms | 1932 | 1948 | 4153 | 1998 | 5977 |
| Ligands | 1 | 0 | 13 | 12 | 10 |
| Water | 222 | 148 | 534 | 330 | 1068 |
| **R.m.s. deviations** | | | | | |
| Bond lengths (Å) | 0.0120 | 0.0112 | 0.0139 | 0.0132 | 0.0164 |
| Bond angles (˚) | 1.6611 | 1.6969 | 1.8883 | 1.7874 | 1.8167 |
| $R_{free}$ (%) | 18.85 | 18.67 | 19.02 | 18.80 | 18.12 |
| $R_{factor}$ (%) | 15.15 | 14.15 | 15.67 | 15.86 | 15.19 |
| Mean B ($Å^2$)[2] | 14.08 | 21.39 | 17.28 | 13.98 | 13.24 |
